# Soluble corn fiber reduces ovalbumin-induced sinonasal inflammation via the gut microbiota-airway axis

**DOI:** 10.1101/2020.07.23.216754

**Authors:** Sierra A. Jaramillo, Emily M. Borsom, Gabrielle M. Orsini, Oliver Kask, Keehoon Lee, Allyson H. Hirsch, Erik Settles, Xiaojian Shi, Haiwei Gu, Andrew T. Koppisch, Nicholas A. Bokulich, J. Gregory Caporaso, Emily K. Cope

## Abstract

Asthma is a chronic airway inflammatory disease that affects approximately 300 million people worldwide, causing a substantial healthcare burden. Although there is a large degree of heterogeneity in the inflammatory response of asthmatics, a subset of patients are characterized by type-2 inflammation, which is in part mediated by T_H_2 cells in both the upper and lower airways. Asthma prevalence is increased in low-socioeconomic-status populations, where disparities in health behavior exist, including a shift toward a western diet characterized by low dietary fiber. Gut microbes metabolize fiber into short chain fatty acids that can reduce type-2 inflammation in peripheral organs, such as the airways. We hypothesized that soluble fiber can reduce ovalbumin (OVA)-induced upper airway inflammation in the context of the unified airway hypothesis, in mice maintained on ingredient-matched western (WD) and control diets (CD) through production of short chain fatty acids. Our results show that soluble fiber reduces *IL-4* and *IL-13* gene expression (p<0.05, Mann Whitney) in the sinonasal cavity of CD-fed mice, but this effect was lost in WD-fed mice. This loss of protection in WD-fed mice parallels compositional changes of the cecal and fecal microbiota. Mice fed a soluble fiber supplement while being maintained on a WD had altered microbial communities characterized by lower abundance of fiber fermentering bacteria. This work can be used to develop effective microbiome-based therapeutics as a low-cost method to reduce asthma morbidity.

**IMPORTANCE:** Previous research has supported that western-style diets, typically high-fat and low-fiber, are associated with changes in the gut microbiome and increased inflammation. Western diets are accessible and prominent in low-socioeconomic-status populations, where asthma rates are highest; however, there has yet to be a low-cost asthma therapeutic. For the first time, we investigated whether supplementation with a physiologically relevant quantity of soluble corn fiber can reduce allergic airway inflammation. Our study supports that soluble corn fiber supplementation is associated with compositional shifts of the gut microbiota and reduced airway inflammation, promoting the use of fiber as a low-cost microbiome modifying therapy to reduce asthma-associated inflammation. However, soluble corn fiber in conjunction with a western diet resulted in an alternate gut microbiome composition and loss of protection against allergic airway inflammation. These findings further support the importance of the gut microbiota in host health.

## INTRODUCTION

Recent studies have demonstrated the myriad of ways that the human microbiota, the collective of microorganisms residing on and in the human body, impacts host health. Recent estimates suggest that there are approximately as many bacterial cells colonizing a host as there are human cells [1,2]. These bacterial cells reside on nearly all body surfaces, with the greatest density of microbiota in the gastrointestinal (GI) tract, and specifically the colon [2]. These host-associated microbiota assist in host metabolism, nutrient acquisition, and immune function [3–5]. Germ-free mice have reduced body weight compared to specific pathogen free mice and are protected against diet-induced obesity, indicating that there is less energy uptake without a gut microbiota [6]. *Clostridium* colonization in the GI tract influences host immunity by promoting extrathymic polarization of anti-inflammatory FOXP3+ regulatory T (Treg) cells via production of short chain fatty acids that inhibit histone deacetylase at the FOXP3 locus [7]. These studies indicate an important role of the GI microbiota and gut health, however, less is known about the role of the GI microbiota in the health of peripheral organs, such as the airways.

The gut microbiome-airway axis has been implicated in asthma [8,9], a complex airway inflammatory disease that causes 10 deaths every day in the United States alone [10]. Asthma is characterized by bronchial hyperresponsiveness and reversible expiratory airflow limitation [11]. Asthma affects approximately 300 million people worldwide [12] and it is estimated that over $54 billion dollars are spent annually on asthma related costs [13]. Incidence of asthma continues to increase; an additional 100 million diagnoses are expected by 2025 [14,15]. There is a large degree of heterogeneity in the asthma inflammatory endotypes. A subset of asthmatics are defined by the T helper 2 (T_H_2) signatures, interleukin (IL)-4, IL-5, and IL-13 [11]. In addition, recent research has supported the unified airway hypothesis in asthma, which posits that asthmatic inflammation is not limited to the lungs, but instead manifests in the upper and lower airways, including the sinuses. Understanding the connections of the human microbiota and upper airway inflammation is vital for understanding the role of the gut microbiota in upper and lower airway disease.

This gut-microbiota airway axis is greatly dependent on fiber fermentation by bacteria residing in the colon [16]. Diet plays a primary role in shaping the gut microbiota [17,18] and it is recognized that the gut microbiota of the western world differs from those in more rural populations, likely due to dietary differences [19,20]. Dietary fiber is a critical component of the diet; according to the American Heart Association, the recommended intake of fiber is 25-35 grams daily. In the US, individuals are consuming an average of 15 grams of fiber per day [21]. Microbial metabolism of dietary fiber results in short chain fatty acids (SCFAs) including propionate, acetate, and butyrate, that can diffuse through the epithelial lining and enter the circulatory system of the host [3,16,22]. SCFAs act on several different immune cells, including those implicated in T_H_2-high asthmatic inflammation [16]. In the bone marrow, short chain fatty acids act on antigen-presenting dendritic cells through the G-protein coupled receptor 43 (GPCR 43) signaling [23–25]. This results in a suppressive dendritic cell phenotype with a reduced capacity to polarize T_H_2 cells, thereby decreasing the T_H_2 inflammatory response, including IL-4, IL-5, and IL-13 [16].

Until recently, asthma has been characterized as an isolated lung disease. However, shared inflammatory responses in the sinonasal cavity and lungs of asthmatics has recently been evaluated; elevated Th2 inflammatory responses in the lungs of asthmatics correlate with the upper airway transcriptome [26]. In addition, the upper airway microbiota is altered in asthmatics and is correlated with host immune phenotype [27,28]. These observations support the existence of the unified airway, which posits that concurrent upper and lower airway disease results from the same pathophysiological process [29]. Previous studies investigating the role of the gut microbiota in asthma have solely focused on the gut microbiota-lung axis and neglected to study the upper airways. Therefore, our understanding of fiber’s influence on the airways needs to be expanded beyond the lungs to the upper respiratory tract.

Our study aims to assess prebiotic fiber as a low-cost asthma therapeutic for individuals in the westernized nations. The reported increase in asthma incidence cannot be solely attributed to genetics; thus, environmental exposures such as diet could be a major contributor. Socioeconomic disparities in health behaviors existing across our population, including increased consumption of a low-cost Western diet that is low in dietary fiber, may contribute to disproportionate asthma rates in low socioeconomic status (SES) communities [30]. Development of dietary interventions or prebiotics, a microbial fermentable ingredient with modulating effects on the gut microbiota and therefore beneficial effects on a host [17], to reduce asthma severity for those in low-SES populations is an exciting avenue of research. We hypothesized that the introduction of a prebiotic fiber supplement will lead to decreased upper airway inflammation in mice consuming a western diet and an ingredient-matched control diet. For the first time, we evaluated the influence of soluble corn fiber (SCF) on the sinonasal cavity in a murine model of allergic airway disease. This preliminary study may pave the way toward development of a gut-microbiome based therapy to reduce airway inflammation.

## RESULTS

To address whether a soluble fiber supplement can reduce allergic airway inflammation in the upper airways of mice maintained on a western or ingredient-matched control diet, the mice were supplemented with a physiologically relevant quantity of soluble corn fiber (Table S1, 0.1305 grams/mouse), equivalent to the recommended 25 grams of fiber intake for the average human. Mice were intranasally exposed to either ovalbumin (allergen) to induce upper airway inflammation or PBS (vehicle). Sinonasal inflammation and sinonasal microbiome, fecal microbiome, and cecum microbiome were assessed.

To determine whether soluble corn fiber (SCF) dietary supplement can reduce upper airway inflammation, we assessed gene expression in the sinonasal cavities of mice in each experimental group (n=62 total, Table 1). Of the 16 genes evaluated, five of the targets were T_H_1 markers, seven were T_H_2 markers, and four were controls (Table S2). Gene expression was calculated for all samples using the ΔΔCt method (Fig. S2). For mice maintained on the control diet (CD) and western diet (CD) mice, we observed increased *IL-4* and *IL-13* sinonasal gene expression in the OVA group compared to the PBS groups. *IL-4* and *IL-13* gene expression were significantly decreased in the CD+OVA+SCF group compared to CD+OVA mice (Fig. 2A, Fig. S3; Mann-Whitney U, *IL-4*, p=0.0205; *IL-13*, p=0.0200). However, there was no significant reduction in *IL-4* nor *IL-13* gene expression of the WD+OVA+SCF group when compared to the WD+OVA group (Fig. 2, Mann Whitney U, p>0.05). We also compared gene expression in SCF supplemented groups across diets and found that *IL-4* trended toward a decrease in the CD+OVA+SCF group compared to the WD+OVA+SCF group (Fig. 2A). However, this decrease in gene expression was not significant (Fig. 2A; Mann-Whitney U, p=0.0693). There were no significant differences between *IL-13* in the CD+OVA+SCF group when compared to the WD+OVA+SCF group (Fig. 2B; Mann-Whitney, p=0.7128). We did not observe any significant differences in T_H_1 markers *IL-1β, IFN-γ, IL-6, TGF-β* (Fig. S2, Mann Whitney U, p>0.05).

**Table 1.** Experimental Groups.

|  | **Control Diet (CD)**<br>**(n=)** | **Western Diet (WD)**<br>**(n=)** |
| --- | --- | --- |
| PBS (vehicle control) | 12 | 12 |
| OVA | 12 | 15 |
| OVA + SCF | 12 | 15 |
| OVA + Cellulose | 8 | 8 |

**Figure 1:**
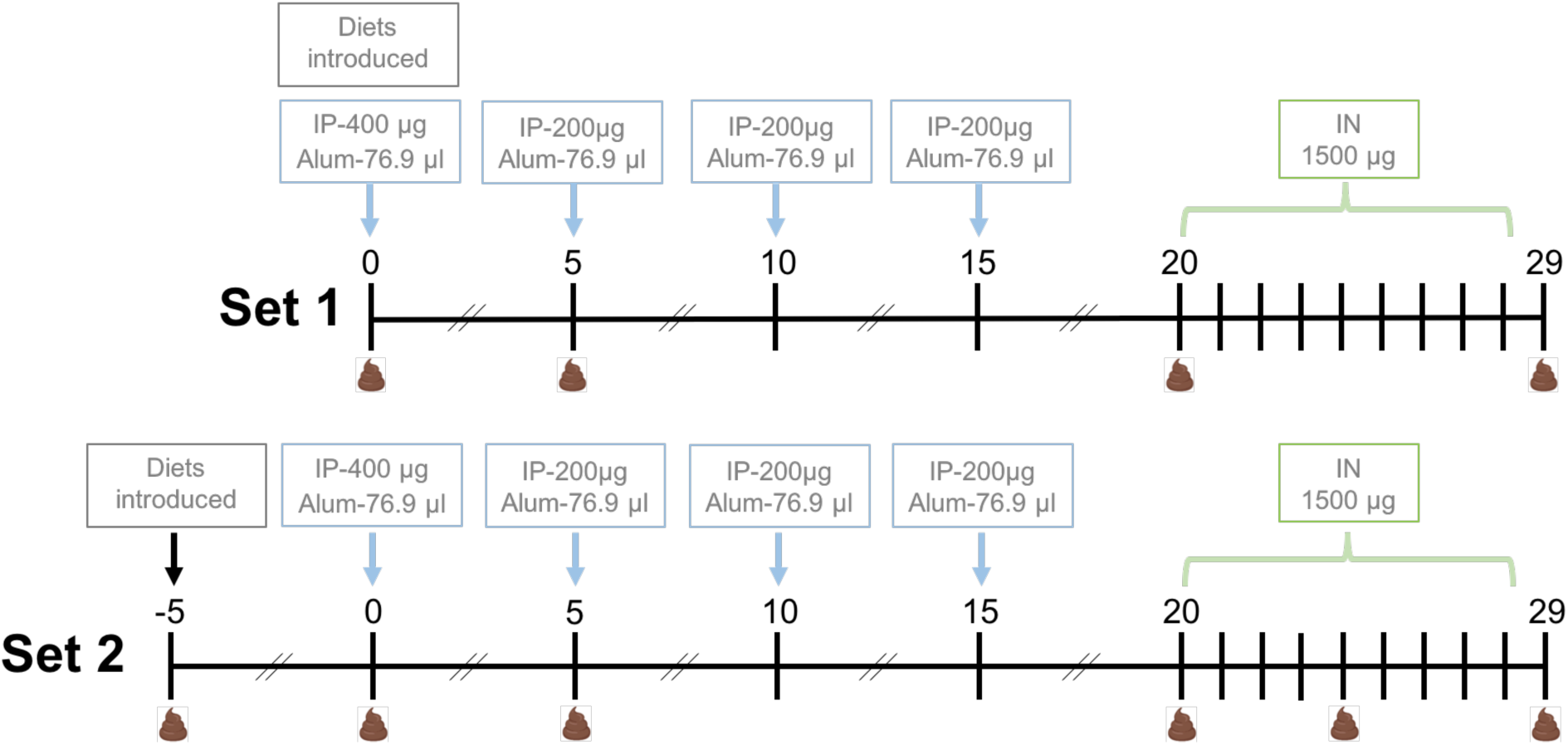
Mouse Timeline Schematic (IP: Intraperitoneal injection with ovalbumin or PBS; Alum: Alum adjuvant added to IP injection, IN: intranasal instillation with ovalbumin or PBS) Set 1 mice were introduced diets on day 0, the same day that intranasal sensitization began. Set 2 mice were introduced to diets 5 days prior to intranasal sensitization. Stool symbols mark timepoints where stool samples were collected.

**Figure 2:**
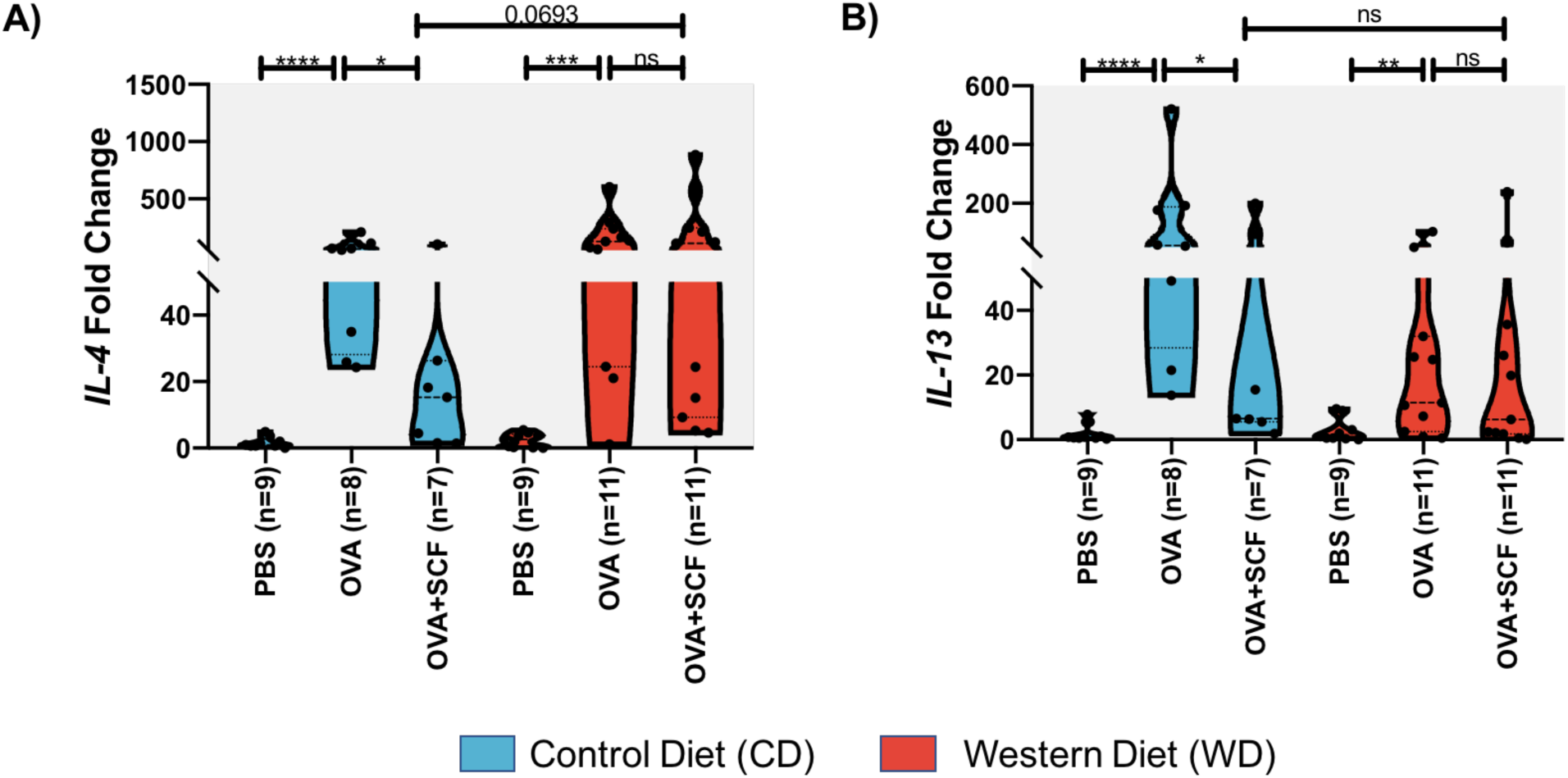
Gene expression of type-2 inflammation markers in mouse sinonasal cavities. Mice underwent allergic airway sensitization with ovalbumin (OVA) or received a vehicle control (PBS). Mice were maintained on either a control diet (CD) or western diet (WD) and a subset were fed a soluble corn fiber supplement (SCF). Reverse transcriptase qPCR was used to measure T_H_2 inflammation markers in the sinonasal cavities. **A)** IL-4, a marker of T_H_2 inflammation was reduced in the sinonasal cavity of fiber treated mice consuming a control diet (Mann-Whitney, p=0.0205) **B)** IL-13, a second marker of T_H_2 inflammation was reduced in the sinonasal cavity of fiber treated mice consuming a control diet (Mann-Whitney, p=0.0200).

Since we observed reduced *IL-4* and *IL-13* gene expression in mice maintained on a control diet (CD) but not mice maintained on a western diet (WD), we evaluated the cecum and fecal microbiota to determine whether changes in the composition or diversity can give insights into our immunological observations. The cecum was included as the site of fiber fermentation in mice, while fecal samples allow non-invasive longitudinal assessments of the gut microbiome. Alpha diversity (within sample) metrics were used to quantify richness, or number of bacterial features present, and Shannon diversity, a metric that weights both the number of features and proportion of each feature, in both the cecum and fecal samples. In the cecum we observed significant inter-diet differences in alpha diversity. Richness (observed Amplicon Sequence Variants [ASVs]) was significantly greater in the CD+OVA, and CD+OVA+SCF groups when compared to the WD+OVA and WD+OVA+SCF groups, respectively (Fig. 3A, Fig. S4, Kruskal-Wallis, FDR corrected, p=0.015, p=0.028). We did not observe significant differences in Shannon diversity in the CD+OVA group compared to the CD+OVA+SCF group or the WD+OVA compared to the WD+OVA+SCF groups (Fig. 3B, Kruskal-Wallis, FDR corrected p=0.421). Richness and Shannon diversity were also measured in terminal fecal samples. We did not observe significant changes in fecal alpha diversity in any comparison for either Richness or Shannon diversity (Fig. 4A, Fig. S4, Kruskal-Wallis, FDR corrected p>0.05), indicating that alpha diversity changes most robustly in the cecum. Volatility plots were used to view alpha diversity of fecal samples throughout the experiment. All treatment groups exhibited a decrease in richness and Shannon diversity within five days of diet implementation. However, the CD+OVA+SCF group maintained greater richness and Shannon diversity than the WD+OVA+SCF group throughout the experiment (Fig. 4B, Fig. 4D). A Linear Mixed Effects model was applied to each alpha diversity metric in the longitudinal samples to determine the effect of diet and supplement (as fixed effects) over time. Results indicate significant differences between diet and supplement groups: SCF-supplemented mice exhibited lower observed richness than cellulose-supplemented mice at baseline (p < 0.001) and less decline in richness over time (p=0.004), though no difference from no-supplement mice. SCF-supplemented mice exhibited lower Shannon H than cellulose-supplemented (p < 0.001) and no-supplement (p=0.004) mice at baseline, and less decline in Shannon H over time compared to cellulose-supplemented mice (p=0.028). Taken together, these results suggest more stable alpha diversity and higher evenness in the SCF-supplemented group.

**Figure 3:**
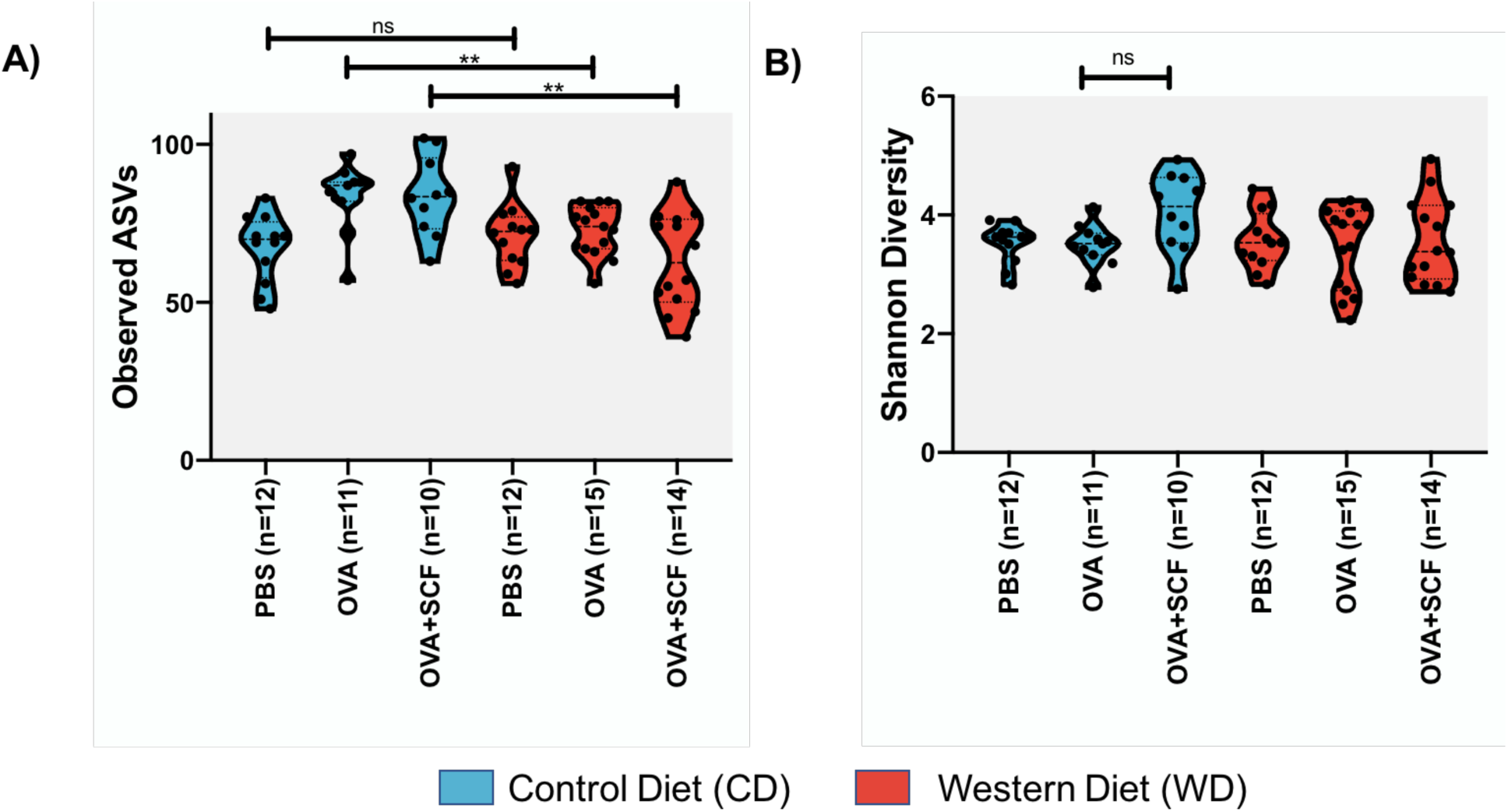
Decreased alpha diversity in cecum of WD treated mice. Mice underwent allergic airway sensitization with ovalbumin (OVA) or received a vehicle control (PBS). Mice were maintained on either a control diet (CD) or western diet (WD) and a subset were fed a soluble corn fiber supplement (SCF). Amplicon sequencing of the V4 16S rRNA gene region was used to assess alpha diversity in the cecum. **A)** Observed amplicon sequencing variants (ASVs) are significantly greater in CD-fed and SCF treated mice than in WD fed and SCF treated mice (p=0.028, Kruskal-Wallis, FDR adjusted p-values) **B)** Shannon diversity is increased in the CD-fed and soluble corn fiber treated group, but it is not significantly different than CD-fed groups without a fiber supplement.

**Figure 4:**
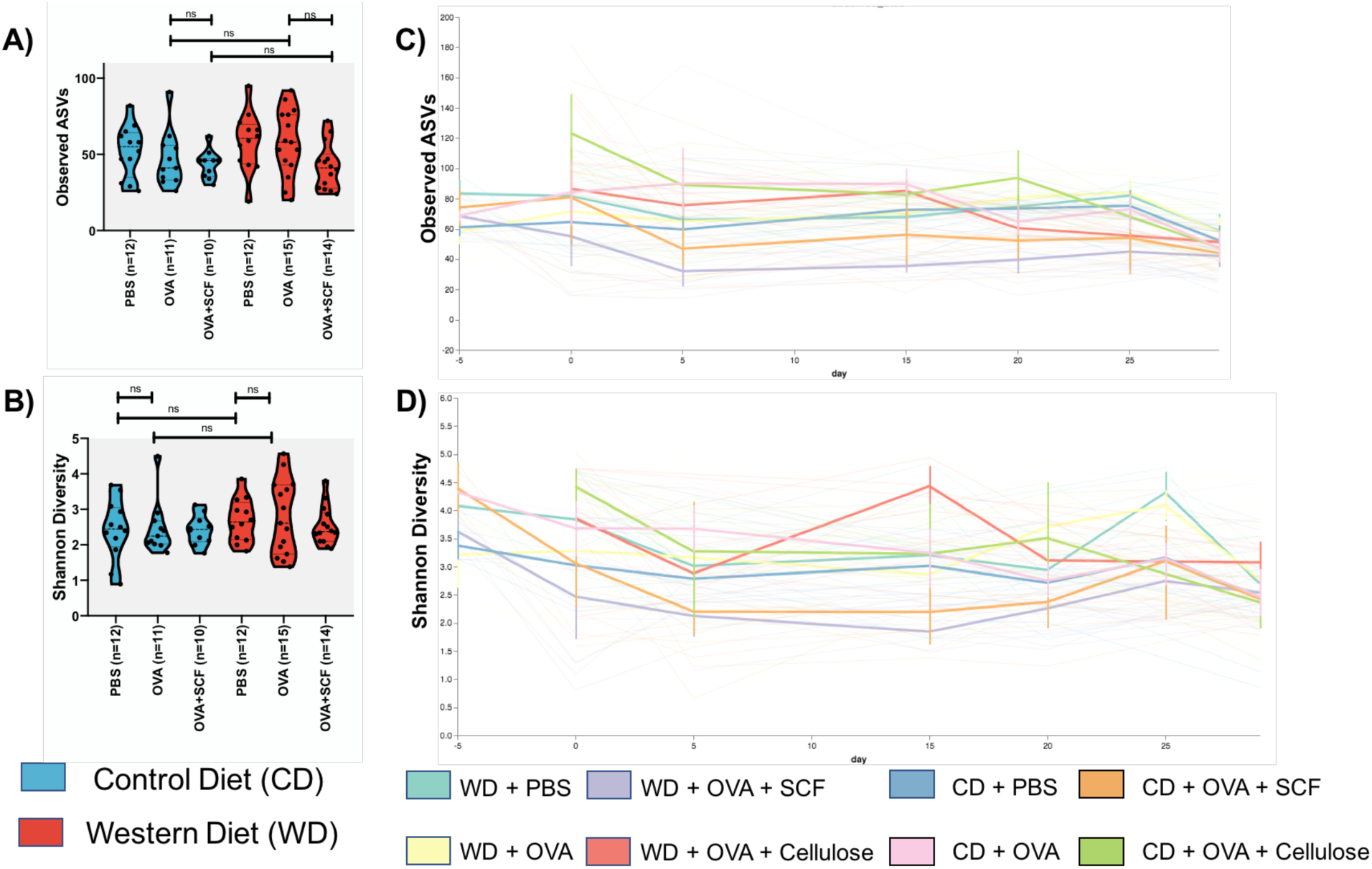
There are no significant differences in alpha diversity of fecal samples (Kruskal-Wallis, FDR adjusted p-values). Mice were maintained on either a control diet (CD) or western diet (WD) and a subset were fed a soluble corn fiber supplement (SCF). Mice were maintained on either a control diet (CD) or western diet (WD). Amplicon sequencing of the V4 16S rRNA gene region was used to assess measures of alpha diversity in the fecal samples collected on the last collection day. **A)** Alpha diversity of fecal samples using the observed amplicon sequencing variants (ASVs) metric **B)** Alpha diversity of fecal samples using the Shannon metric shows no significant differences **C)** Observed ASVs of fecal samples throughout the experiment **D)** Shannon diversity of fecal samples throughout the experiment.

Beta diversity (between-sample) metrics were used to detect compositional changes in the bacterial microbiome between samples. In the cecum, Jaccard, Bray-Curtis, Unweighted Unifrac, and Weighted Unifrac were used to assess beta diversity (Table S3). Principle coordinate analysis (PCoA) of Jaccard and Bray-Curtis metrics demonstrate significant alterations in the the cecum microbiota based on diet and SCF supplementation (Fig. 5, PERMANOVA Jaccard diet: p=0.001, supplement: p=0.001; Bray-Curtis diet: p=0.001, supplement: p=0.001). A Random Forest machine learning sample classifier trained on the cecum microbiota revealed which taxa were most important in predicting SCF supplement status. Genera ranked as high importance of predicting SCF supplementation in the cecal microbiome included *Ruminococcus, Coprococcus, Coprobacillus, Blautia*, and *Clostridium* (Fig. 6, overall accuracy : 0.944444, AUC SCF: 1, AUC cellulose : 0.666667, AUC no supplement: 1). Many of these taxa are known short chain fatty acid producers and have been associated with gastrointestinal health.

**Figure 5:**
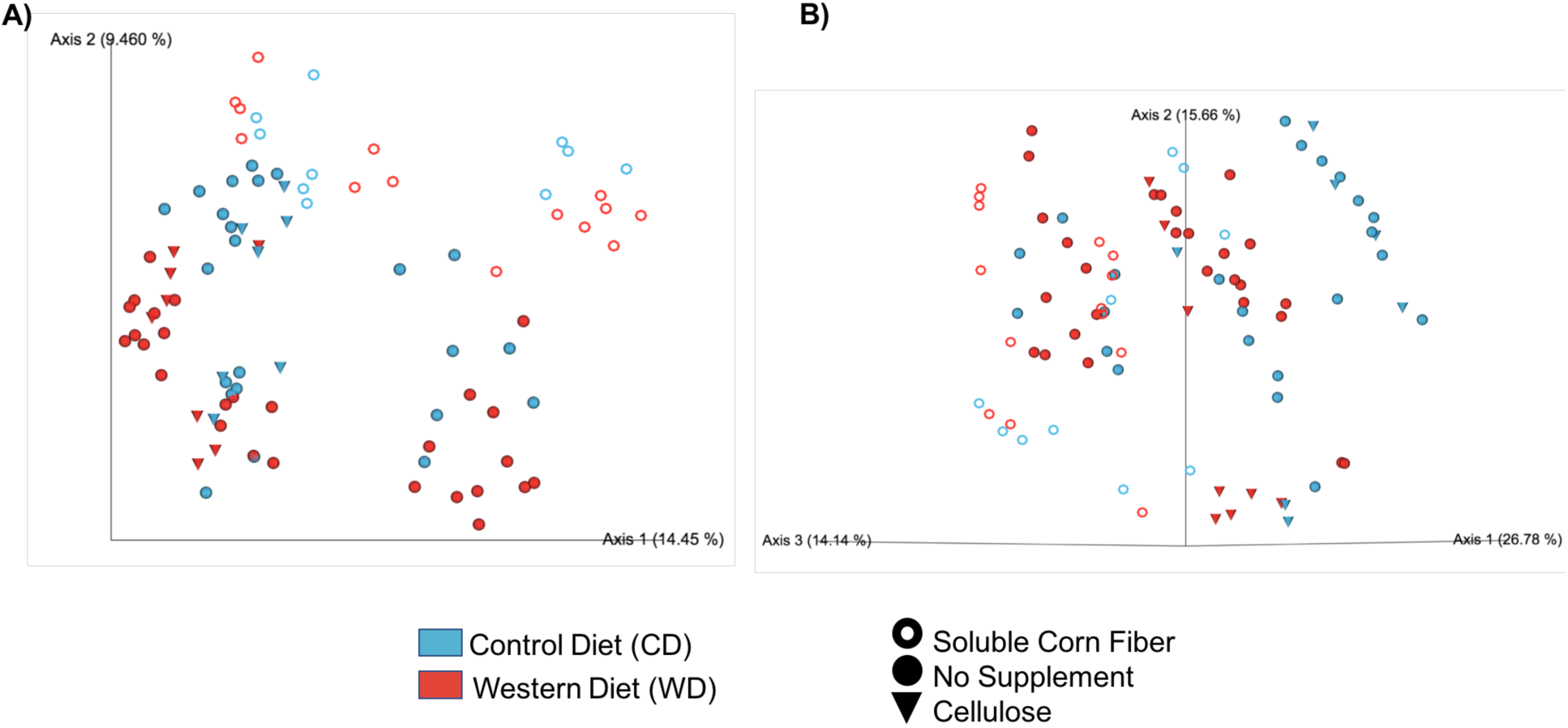
SCF treated mice have distinct cecal microbiomes. Mice underwent allergic airway sensitization with ovalbumin (OVA) or received a vehicle control (PBS). Mice were maintained on either a control diet (CD) or western diet (WD). Mice were supplemented with either soluble corn fiber (SCF), an insoluble fiber (Cellulose), or received no supplement. Amplicon sequencing of the V4 16S rRNA gene region was used to assess measures of beta diversity in the cecum samples collected on the last collection day. **A)** A PCoA of a Jaccard distances indicates that diet and supplement are significant variables in the cecal microbiomes (diet: PERMANOVA p=0.002, pseudo-f=3.65169; supplement : p=0.001, pseudo-f=5.5741). **B)** A PCoA of Bray-Curtis distances indicates that diet and supplement are significant variables in the cecal microbiomes (diet: p=0.001, pseudo-f=6.292; supplement: p=0.001, pseudo-f=8.6421).

**Figure 6:**
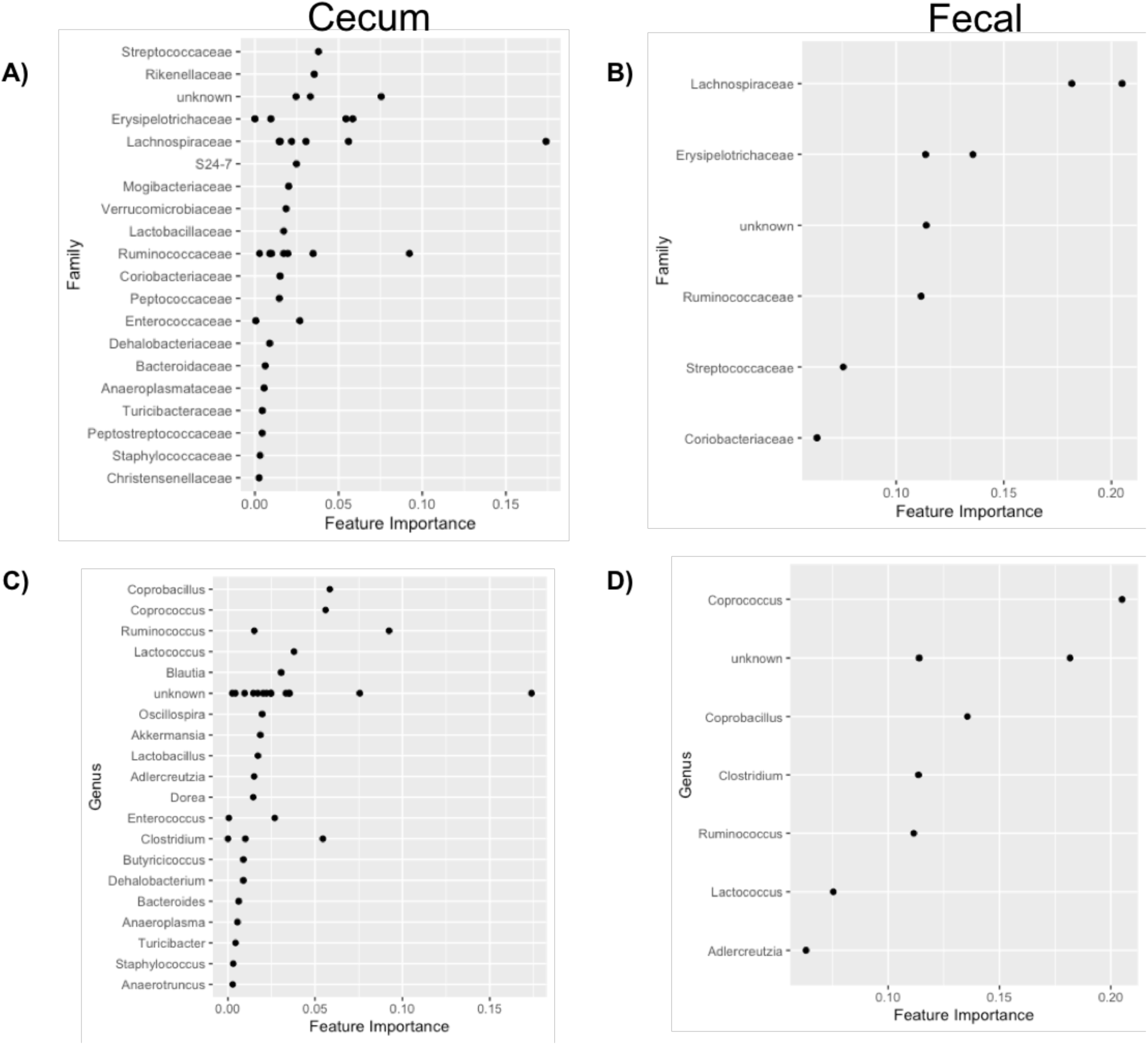
Random Forest feature selection identifies fecal and cecal bacterial genera and families associated with diet and supplementation. Mice underwent allergic airway sensitization with ovalbumin (OVA) or received a vehicle control (PBS). Mice were maintained on either a control diet (CD) or western diet (WD) and a subset were fed a soluble corn fiber supplement (SCF), insoluble fiber (Cellulose), or received no supplement. Amplicon sequencing of the V4 16S rRNA gene region was performed. Random Forests was used to determine which taxa were most important in predicting cecal microbiome composition. Gut microbiota accurately predicts diet and supplement status. Cecal microbiome overall accuracy: 0.944444, AUC SCF: 1, AUC cellulose: 0.666667, AUC no supplement : 1 **A)** Importance of family to predict supplement effect on cecal microbiome **B)** Importance of family to predict supplement effect on fecal microbiome **C)** Importance of genus to predict supplement effect on cecal microbiome Fecal microbiome overall accuracy: 0.888889, AUC SCF: 1, AUC cellulose: 0.333333, AUC no supplement: 1 **D)** Importance of genus to predict supplement effect on fecal microbiome.

SCF supplementation did not result in reduced *IL-4* or *IL-13* gene expression in mice maintained on a WD. Therefore, we hypothesized that the cecum microbiome in WD-fed mice had a distinct composition characterized by reduced abundance of fiber fermenting microbiota. To test this hypothesis, we examined the cecum microbiome between the CD+OVA+SCF and WD+OVA+SCF mice using Jaccard, Bray-Curtis, Unweighted Unifrac, and Weighted Unifrac diversity metrics (Table S3). A PCoA of Jaccard demonstrated that the CD+OVA+SCF microbiomes clustered separately than the WD+OVA+SCF group (Fig. 7A, PERMANOVA p=0.017). Analysis of composition of microbiomes (ANCOM) was performed to determine which taxa are differentially abundant between the two groups. One taxon, *Rikenellaceae*, was enriched in the CD+OVA+SCF mice, and was absent in the WD+OVA+SCF group (Figure 7B, ANCOM, W=41).

**Figure 7:**
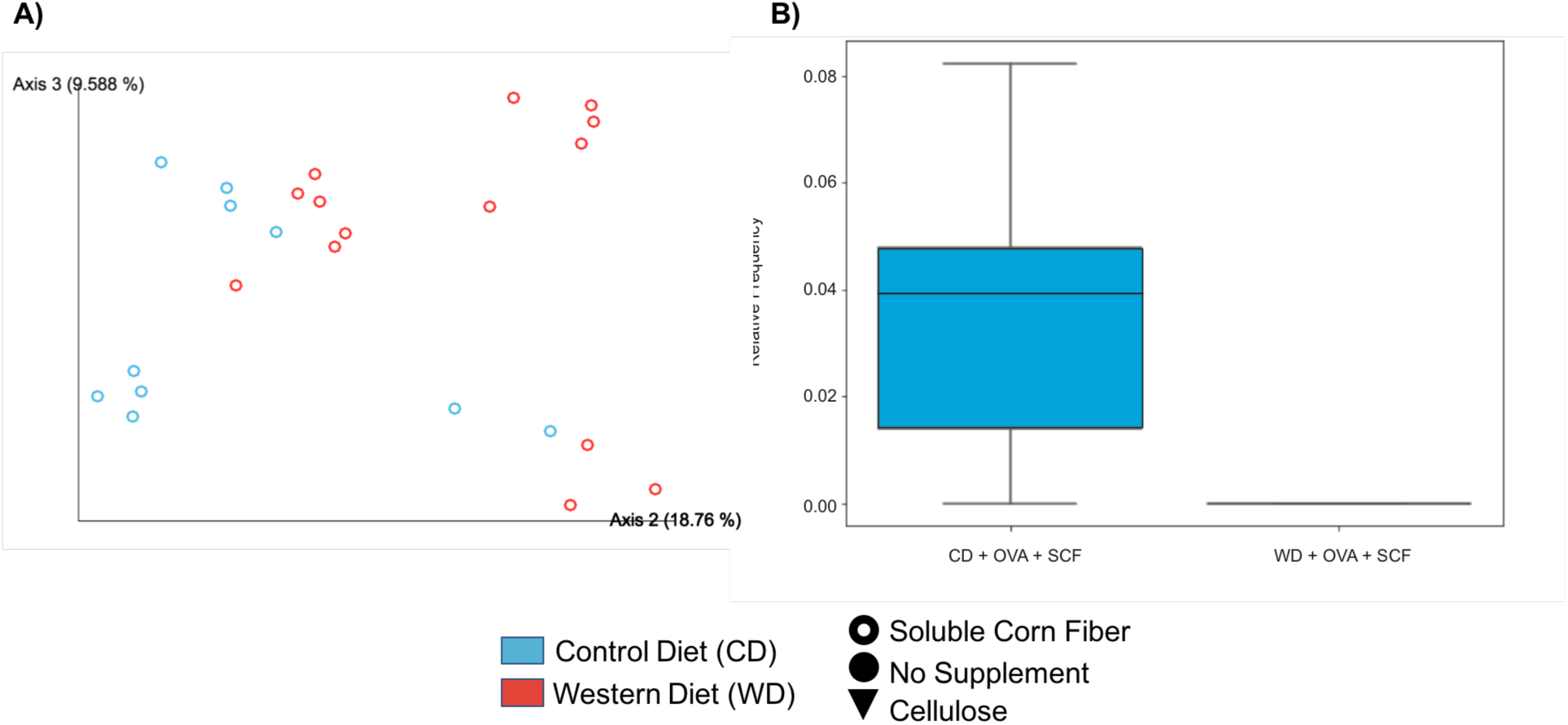
SCF treated mice have diet caused cecal microbiome differences. Mice underwent allergic airway sensitization with ovalbumin (OVA) and received a soluble corn fiber supplement (SCF) while being maintained on either a control diet (CD) or western diet (WD). Amplicon sequencing of the V4 16S rRNA gene region was performed. Beta diversity was used to assess diet based differences in the SCF treated mice. **A)** Multivariate permutation testing of PERMANOVA based on Jaccard distanced shows that diet and supplement are significant variables in the cecal microbiomes (diet:p=0.011, pseudo-f=2.40083) **B)** Relative frequency of *Rikenellaceae* in the cecal microbiome. SCF treated mice consuming a CD have increased relative frequency of *Rikenellaceae* compared to SCF treated mice consuming a WD (ANCOM, W=41).

Prior studies have shown that diet is a major driver of gut microbiome composition [18,31], and in this study, we observe diet-specific differences in host immune response. To address longitudinal diet-induced changes in the microbiota, fecal samples were collected immediately before diet implementation (baseline samples) and throughout the experiment. Beta diversity measures were used to assess the effect that diet changes had on the fecal microbiota (Table S3). A PCoA of the Jaccard metric shows that the baseline fecal samples are compositionally distinct from samples collected on all other days throughout the experiment (Fig. 8A, PERMANOVA p=0.001). After 5 days of consuming the new experimental diets there is a robust shift in the fecal microbiome. To determine how SCF modified the fecal microbiome over time, we plotted a PCoA of the Jaccard distance metric using time as a custom horizontal axis (Fig. 8B). These results show that all baseline fecal samples are clustered together, as would be expected since the mice were not yet introduced to diet or SCF. However, after 5 days on the diet and SCF, SC-treated mice cluster separately from all other fecal samples (Fig. 8B). These results demonstrate a rapid and robust change in the fecal microbiota after just 5 days on a prebiotic SCF-supplement. LME was performed on the first principle coordinate axis (PC1) from PCoAs generated from each beta diversity metric to determine the effect of diet and supplement (as fixed effects) using time as a random effect. These results indicate significant differences between diet and supplement groups. Using weighted and unweighted metrics, the bacterial microbiota of SCF-supplemented mice were distinct from cellulose-supplemented (p < 0.001 Jaccard, p < 0.001 Bray Curtis, p < 0.001 unweighted UniFrac, and p=0.001 weighted UniFrac) and non-supplemented mice (p=0.002 Jaccard, p=0.001 Bray Curtis, p=0.015 unweighted UniFrac, and p=0.001 weighted UniFrac) at baseline, when diets were administered. There were no significant differences over time by diet alone, however diet did interact with supplement using weighted metrics (p=0.039 Bray Curtis, p=0.008 Weighted UniFrac). Also using weighted metric, SCF-fed mice were distinct from cellulose (p=0.023 Weighted UniFrac) and non-supplemented mice (p < 0.001 Weighted UniFrac, p=0.024 Bray Curtis) over time. These results support that diet interacts with SCF to alter the fecal microbiome over time. The bacterial microbiota of SCF-supplemented mice on the control diet were distinct from non-supplemented (p=0.026 Jaccard) mice and cellulose-supplemented mice (p=0.047 Jaccard) maintained on the Western Diet.

**Figure 8:**
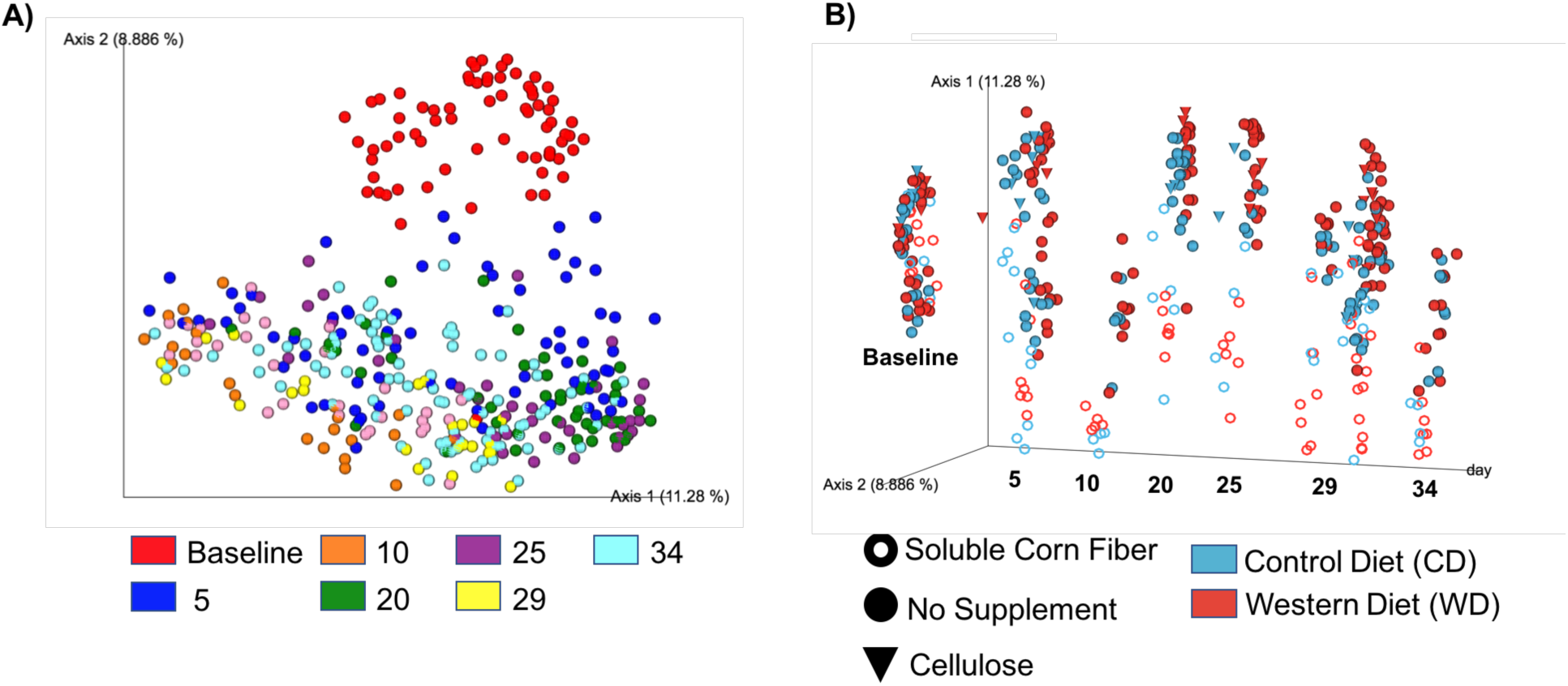
Mice were introduced to a control diet (CD) or western diet (WD) with either no supplement, a soluble corn fiber (SCF) supplement, or an insoluble fiber (Cellulose) supplement day 0 or day -5 (baseline). Fecal samples were collected throughout the experiment and used to assess gut microbiome changes. Microbiome changes were determined by sequencing the 16S V4 rRNA gene region. Five days after diet and fiber supplementation a distinct fecal microbiome is developed. **A)** The fecal microbiome shifts by day 5 of the experiment, but little changes in the feces occur after five days of diet and supplement changes (Jaccard PERMANOVA p=0.001, pseudo-f=33.4822). **B)** Using time as a custom axis, it is shown that the SCF treated mice develop distinct fecal microbiomes after five days of diet and supplementation.

Finally, we assessed the bacterial microbiota of the sinonasal tract to determine whether SCF supplementation or alterations in the sinonasal immune response changed the composition or diversity of these upper airway microbial communities. There were no differences in Richness and Shannon diversity of sinonasal samples (Fig. S5, Kruskal Wallis, FDR corrected p>0.05). Nor did we observe a significant compositional shift in Jaccard, Bray-Curtis, Unweighted Unifrac, and Weighted Unifrac metrics based on diet or supplement (Table S5, PERMANOVA, p>0.05).

We hypothesized that circulating short chain fatty acids act on peripheral immune populations to reduce airway inflammation. To test this, we measured serum short chain fatty acids, including acetic acid, butyric acid, and propionic acid, were via gas chromatography-mass spectrometry (GC-MS). We did not detect changes in serum short chain fatty acid levels between groups (Fig. 9). Other less common short chain fatty acids were also detected. These other short chain fatty acids also showed little variation between groups (Fig. S6, Mann-Whitney U, p>0.05). Histological analysis Periodic Acid Schiff (PAS)-stained of sinonasal sections revealed non-significant changes in goblet cells in the upper airways. Goblet cells increased in OVA treated mice and we observed a slight, but non-significant decrease in OVA+SCF mice in both diet groups, and a non-significant increase in goblet cell counts in the WD treated groups than in the CD treated groups (Fig. S7, Mann-Whitney U, p>0.05).

**Figure 9:**
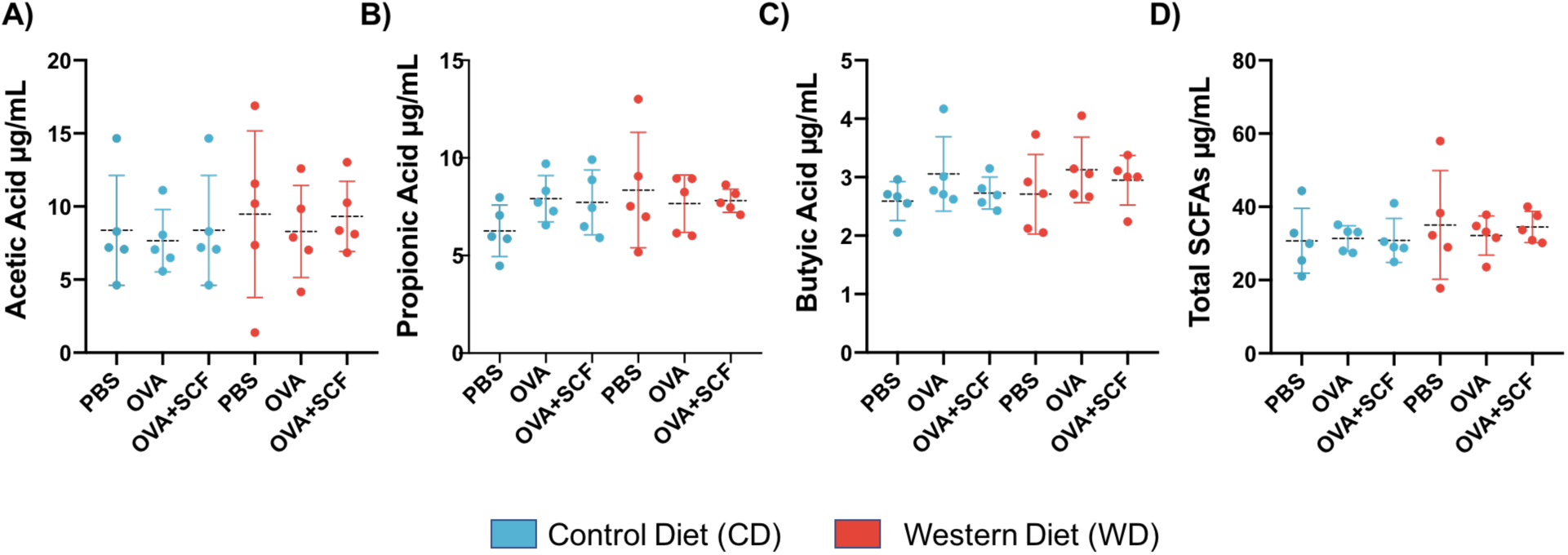
Mice were maintained on a control diet (CD) or western diet (WD) with either no supplement, a soluble corn fiber (SCF) supplement, or an insoluble fiber (Cellulose). Serum was collected at the end of the experiment. Gas chromatography-mass spectrometry was used to quantify short chain fatty acids in serum samples. (CD: control diet, WD: western diet, PBS : no allergic airway sensitization, OVA : allergic airway sensitization with ovalbumin, OVA + SCF : allergic airway sensitization and a soluble corn fiber supplement). Mann-Whitney U tests showed no significant differences for **A)** Acetic acid **B)** Propionic acid **C)** Butyric acid and **D)** Total short chain fatty acids.

## DISCUSSION

Previous research has supported the use of fiber to reduce allergic airway inflammation in the lungs [16], but has not been aimed at its effect on the unified airways. Our study aimed to understand the effect of dietary fiber on allergic inflammation in the sinuses. We sensitized mice with ovalbumin and measured expression of T_H_2 cytokines, including *IL-4* and *IL-13. IL-4* and *IL-13* promote smooth muscle constriction, eosinophil recruitment, immunoglobulin E class switching, and differentiation of naive CD4+ T cells into T_H_2 cells. We found that CD-fed mice consuming a SCF supplement had reduced levels of *IL-4* and *IL-13* gene expression in the sinonasal cavity. This demonstrates, for the first time, that the anti-inflammatory effects of a soluble fiber supplement are not limited to the lower airways, but also affect the upper airways. These results further support the use of soluble fiber as a prebiotic treatment for allergic airway inflammation. Future studies will focus on the role of soluble corn fiber in upper airway inflammation associated with asthma or chronic rhinosinusitis (CRS), a common airway inflammatory disease of the sinonasal tract.

Prior studies of soluble fiber and respiratory disease have used a poorly fermentable fiber, cellulose, to control for differences in mineral uptake [16,32]. In humans, insoluble fiber contributes to stool bulk, whereas soluble fiber is readily fermented by gut microorganisms into anti-inflammatory short chain fatty acids [33,34]. In our study, the CD-fed cellulose-treated mice also had reduced sinonasal *IL-4* and *IL-13* gene expression (Fig. S3, Mann Whitney, p=0.0031), illustrating that insoluble fiber can also reduce airway inflammation. This finding aligns with a recent study that observed decreases in airway inflammation markers in mice fed both pectin (a readily fermentable fiber) and cellulose (an insoluble fiber, [35]). We hypothesize that this result may be due to anatomical, physiological, and compositional differences in the gut microbiota between mice and humans [36]. We may be observing limited cellulose fermentation of insoluble fiber in the mouse GI tract or other trophic effects of insoluble fiber independent of microbial fermentation [37]. It is important to note that further investigation into a combination of soluble and insoluble fiber may be useful to study. There is potential for cellulose to contribute to digestion beyond stool bulk even in humans.

To determine if soluble fiber can be used as a microbiota-modifying therapy with clinical benefits for asthmatics in industrialized nations, we implemented an ingredient matched western style diet into our study; the only dietary differences in our study were percentages of carbohydrates and fat. The WD-fed group was treated identically to the previously described CD-fed mice. However, in the WD-fed group, *IL-4* and *IL-13* were not reduced by SCF supplementation, whereas SCF supplementation resulted in reduction of *IL-4* and *IL-13* gene expression in the CD-fed group. Previous research has supported that high-fat diets are associated with increased airway inflammation, as seen by increased neutrophil counts in the lungs after consuming a high-fat meal in humans [38]. In one study, a high fat diet was associated with impaired airway dendritic cell function and reduced regulatory T cells [39]. We originally hypothesized that short chain fatty acids act on dendritic cells to promote a suppressive phenotype, it’s possible that these effects are blunted if dendritic cells have an altered phenotype due to diet alone. In mice, high fat diets are associated with alterations in the gut microbiota ncharacterized by increased proportions of Firmicutes, including *Molliculites* and *Ruminococcaceae* and decreased proportions of Bacteroidetes including *Rikenellaceae* [31,40]. We also observed a depletion in richness and a shift in the microbial composition in WD+SCF-fed mice relative to mice in the CD+SCF-fed group, indicating that there is a loss of taxa at the site of fiber fermentation in the WD-fed group. We hypothesize that the decreased ASVs observed in the WD+OVA+SCF mice of our study are a result of a loss of fiber fermenting bacteria. Our results suggest that fiber supplementation alone, without improvement of other aspects of diet or inoculation of a beneficial fiber fermenting species, may not be sufficient to reduce allergic airway inflammation.

We used a Random Forest machine learning classifier to identify which taxa are important in predicting the diet and supplement using cecal microbiota composition. *Ruminococcus, Coprococcus, Coprobacillus, Blautia*, and *Clostridium* were ranked highest in importance for differentiating diet and SCF status. *Ruminococcus, Blautia* and *Clostridium* have previously been identified as fiber fermenting organisms, with increased prevalence in individuals consuming fiber rich foods [3,41–43]. We hypothesize that the taxa lost in the WD-fed group were these fiber fermenting microbes. To investigate this further, we compared the relative abundances of taxa between the WD+OVA+SCF group and the CD+OVA+SCF group. Notably, the family *Rikenellaceae* was present in the cecal samples of CD+OVA+SCF mice, but absent from the WD+OVA+SCF mice. *Rikenellaceae* has been correlated with a high fiber diet where it has been considered protective against forms of colitis [44]. Future research will focus on whether members of *Rikenellaceae* are critical for fiber-mediated reduction of allergic airway inflammation. Future studies will address whether administering members of the *Rikenellaceae* family as a probiotic in conjunction with SCF (a synbiotic) may reduce T_H_2 inflammation even in a WD-fed group.

Fecal samples are often used as a non-invasive way to characterize microbiota composition and diversity in a variety of mammalian systems. We investigated whether the compositional differences that we observed in the cecum, the site of fiber fermentation in mice, could be detected in the fecal samples. We observed compositional shifts in the fecal microbiota driven by SCF supplementation, but a loss of signal in our alpha diversity metrics in the fecal microbiome compared to the cecum microbiome. Because these were fecal samples it is unknown whether these compositional shifts are transient, e.g. from the cecum, or if this is reflective of shifts in the mucosal colonic microbiota. Notably, performing 16S rRNA gene sequencing of fecal samples is customary and instrumental in human and murine studies. Our findings add further support that fecal microbiome analysis is necessary, but may not be reflective of the entirety of changes in along intestinal sites. This further supports the need to carefully choose sampling sites during microbiome studies.

To determine if the decreased T_H_2 inflammation in the CD-fed and SCF supplemented group can be attributed to SCFA production, we measured circulatory short chain fatty acids. We observed no difference in acetic acid, butyric acid, propionic acid, or total short chain fatty acids. This result differs from previous research and the current hypothesized mechanism for short chain fatty acids as a metabolite able to reduce T_H_2 inflammation. This finding may be due to several factors, including our relatively low, physiologically relevant dose of fiber consumed in this study, relative to other published murine studies that supplement 30% of their diet with fiber [16,45]. Increasing the sample size may have resulted in increased statistical power to detect changes. However, other mechanisms may influence fiber’s effect on T_H_2 inflammation. Dendritic cell activation and T_H_2 cell differentiation is dependent on many factors, including transcription factors such as GATA3 and STAT6 [46,47], thymic stromal lymphopoietin (TSLP) [48], dose dependencies of lipopolysaccharide (LPS) [49], and activation of pattern recognition receptors [50]. Perhaps the altered gut microbiota in SCF-treated mice can mitigate dendritic cell activation and T_H_2 cell polarization through one of these other mechanisms. Indeed, fiber fermenting microbes possess exogenous or endogenous signals that decrease dendritic cell activation [16,51]. There is a need for further research into potentially new host-microbiota interactions mediated by fiber supplementation. In future studies, we will evaluate the total metabolome in the cecum, fecal, and serum.

We demonstrate reduced T_H_2 inflammation in an OVA-induced model of allergic airway inflammation. Although OVA is a widely used antigen to induce IgE-mediated allergic reactions in mice, future studies in our lab will include other relevant allergens, including cockroach allergen and fungal extract, to address whether the effect we observed is translatable across common allergy models and distinct strains of mice. In this study, we quantified short chain fatty acids in the serum of mice, but future studies will include SCFA quantification in the cecum, feces, and upper airways. Our study did not identify the mechanism by which microbial fermentation of soluble fiber reduced T_H_2 inflammation in the upper airways. We hypothesize that the mechanism is similar to what was reported by Trompette and colleagues [16], but future work will include unbiased metabolomics and use of GPR41 and GPR43 knockout mice to determine whether microbial derived short chain fatty acids signal through G protein coupled receptors to alter dendritic cell function in the upper airways. Finally, we did not assess the phenotype of specific cell types contributing to allergic inflammation, including dendritic cells, CD4+ T_H_2 cells or IL-13 producing type 2 Innate Lymphoid Cells (ILC2s). Our study was not set up to determine the cellular origin of the type-2 cytokines; future studies will include a more detailed evaluation of contributing airway immune cells.

In conclusion, our study aimed at testing a low-cost prebiotic fiber supplement as an airway inflammation therapeutic in western nations, where fiber consumption is reduced. These results highlight the role of the gut microbiota-airway axis in upper airway disease, for which protective gut organisms are necessary for health benefits. Our study is unique in that it evaluates the role of soluble fiber on the gut microbiome and upper airway disease in mice maintained on ingredient matched Western and Control diets. SCF was shown, for the first time, to have anti-inflammatory effects on upper airways of control diet-fed mice, but this effect was lost in mice fed a western diet. We hypothesize that we will observe anti-inflammatory effects of SCF in a human population consuming a western diet, despite our findings in mice, because humans will consume a more varied diet over time. We hope that this research can be expanded on to reduce disease incidence of asthma in all populations, including vulnerable low socioeconomic-status populations.

## MATERIALS AND METHODS

### Murine Model

All mouse experiments were approved by the International Animal Care and Use Committee at Northern Arizona University (IACUC approval number 16-008). Female Balb/c mice (Jackson Laboratory) at 7 weeks old were randomly separated into 2 diet groups: 32 mice in the high-fat western diet (WD) group and 32 mice in the low-fat control diet (CD) group. Mice in each diet group were randomly split into 4 subgroups each containing 8 mice. The 4 subgroups are: 1) mice that received a vehicle control of PBS without a fiber supplement (PBS) 2) mice with induced allergic airway inflammation (OVA) 3) mice with induced allergic airway inflammation and a soluble corn fiber supplement (OVA + SCF) 4) mice with induced allergic airway inflammation and a cellulose, insoluble fiber, supplement (OVA + cellulose). All mice described previously began their diets and allergic airway sensitization on the same day. A second set of animal experiments was conducted for which mice began their diets five days prior to allergic airway sensitization (Fig. 1). The replicated experiment remained the same as previously written, but with the removal of the cellulose group. Four mice were included in each group with the exception of the WD + OVA and WD + OVA + SCF groups which had 7 mice each (Table1). The sample size of the WD+OVA and WD+OVA+SCF group were increased because they were the variable groups used to test our hypothesis.

### Diets

All diets were designed to match the typical American western style diet. Diets and supplements were provided by cage. The macronutrients and supplement amounts are as follows. The high-fat western diet consisted of 40.3% Kcal fat,18.2 % Kcal protein, and 41.5% Kcal carbohydrates (Envigo, Teklad custom diet: TD.180420; Fig. S1). The ingredient matched control diet consisted of the same ingredients, but at macronutrients of 12.6% Kcal fat, 18.1% Kcal protein, and 69.3% Kcal carbohydrates (Envigo, Teklad custom diet: TD.180421) (Table 2). Due to cellulose being an insoluble fiber, 2 additional diets were developed to supplement mice with approximately the same volume of 0.1305 g of cellulose per day (Envigo, Teklad custom diets: TD.180417,TD.180418; Supplemental Table 1). The soluble corn fiber (SCF) supplement was supplied by Tate and Lyle (Promitor85). This fiber was chosen because it has been well tolerated in human studies [52]. SCF was added to the drinking water at 0.1305 grams SCF per 3 milliliters of water. This volume was determined based on an equation from Hintze and colleagues as follows(*unit Nutrient/ day*)*/*(2070 *kcal/day*) *∗* (4.4 *kcal/g diet*) *∗* 1000 = (*unit Nutrient/kg diet*) [53] in addition to understanding that 3 milliliters of water is the average water intake of a 15 gram body weight Balb/c mouse per day [54].

**Table 2.** Diet Formulas: No Supplement Added

| **Formula** | **Control Diet (g/kg)** | **Western Diet (g/kg)** |
| --- | --- | --- |
| Casein | 190 | 236 |
| DL-Methionine | 3 | 3.54 |
| Sucrose | 50 | 179.12 |
| Corn Starch | 520.59 | 189.2 |
| Maltodextrin | 120 | 120 |
| Anhydrous Milkfat | 19.5 | 100 |
| Beef Tallow | 19.5 | 100 |
| Soybean Oil | 11 | 3.5 |
| Cellulose | 17.4 | 10.8 |
| Mineral Mix | 35 | 41.3 |
| Calcium Phosphate | 4 | 4.72 |
| Vitamin Mix | 10 | 11.8 |
| Ethoxyquin | 0.01 | 0.02 |

### Allergic Airway Sensitization

All OVA solutions were made in 103.1 µl of phosphate-buffered saline (PBS) then filter sterilized in a 0.22 µm filter. Filter sterilized OVA + PBS was added to 76.9 µl of sterile alum adjuvant (Thermo Fisher) per mouse. OVA intranasal installation solutions were made by combining 1500 µg of OVA per 40 µl of PBS then filter sterilized. Each group of mice received either allergic airway inflammation using ovalbumin (OVA) or a vehicle control (PBS) following modified methods from Mendiola et al [55]. In short, mice were sensitized using intraperitoneal (IP) injections on days 0, 5, 10, and 15. The day 0 IP injection contained 400 µg of OVA and 76.9 µL of alum adjuvant in 103.1 µl of sterile phosphate buffered saline solution (PBS) (Sigma-aldrich). Mice received IP with 200 µg OVA + 76.9 µL of alum adjuvant 103.1 µl of sterile PBS on days 5, 10, and 15. Control mice received 180 µl PBS via IP injection at each timepoint. Mice received intranasal installations of either 1500 µg OVA in 40 µl of PBS or the vehicle control of 40 µl PBS each day from day 20 through 29. On intranasal installation days each mouse received a 20 µl intranasal installation, had a 30 minute period of rest then received a second 20 µl intranasal installation. These methods were modified by adding an additional 20 µl of PBS per installation to better sensitize the lower airways in addition to the upper airways.

### Sample Collections

Fecal samples were collected on days 0, 5, 10, 15, and 20 through 29 for a longitudinal gut microbiome analysis (Fig. 1). Fresh fecal samples were immediately placed at -70°C. On day 29, mice were euthanized using inhaled carbon-dioxide. Blood was collected via cardiac puncture and placed in a plastic 1.5 mL tube to be used for GC-MS. Cecum and sinonasal cavities were collected from all mice and immediately placed in RNAlater (Qiagen). Samples in RNAlater were placed at 4°C for 24 hours and then stored at -70°C. Sinonasal samples for 2 mice per group were dissected and immediately placed in Carnoy’s solution for histological analysis.

### DNA/RNA Extractions

DNA and RNA were extracted in parallel using the Qiagen AllPrep Kit (Qiagen) with an additional mechanical lysis. Briefly, samples were placed in a lysing matrix E tube (MP Biomedical) with 600 µl of Buffer RLT Plus and lysed in 30 second increments for a total of 6 minutes at 10,000 x g. Samples were sat resting for 30 seconds between each bead beating to prevent heating. Extraction continued following the manufacturer’s protocol. Both DNA and RNA were quantified using a NanoDrop 2000. Quantified DNA from cecum, fecal, and sinonasal samples were used for 16S rRNA targeted gene sequencing and sinonasal RNA was used for cytokine detection. Samples were extracted in random sets of 94 samples to eliminate extraction bias. Every set of 94 samples had an additional 2 extraction blank controls. Extraction blanks did not contain any tissue during the extraction, but were carried throughout the entire extraction and 16S rRNA gene sequencing.

### Airway Inflammatory Gene Expression

Genomic DNA was removed from RNA using the Rapid-Out DNA removal-kit (ThermoFisher). Reverse transcription was performed using the High-Capacity cDNA Reverse Transcription Kit (ThermoFisher) and complementary DNA (cDNA) was quantified using the NanoDrop 2000. A custom RT2-qPCR array (Qiagen) was designed to detect asthma inflammatory profiles along with the endogenous control glyceraldehyde 3-phosphate dehydrogenase (GAPDH). Four T_H_1 genes and seven T_H_2 genes were targeted on the custom array (Supplemental Table 2). Additional quality controls consisting of a genomic DNA control, reverse transcription control, and positive PCR control were included for every sample. One nanogram of cDNA was added to the custom RT2-array and screened on the QuantStudio 12k flex system (Applied Biosystems) with the following PCR conditions, 95°C for 10 minutes for Taq activation and 40 cycles of 95°C for 15 seconds and 60°C for 1 minute. Data was analyzed following the delta-delta Ct (DDCt) method using GAPDH as the endogenous control. During analysis, CD + PBS mice were considered controls for all other mice on the control diet and WD + PBS mice were considered controls for all mice on the western diet.

### Histology

To examine histological sections of sinonasal cavities, mice were decapitated and mouse heads were immediately placed in Carnoy’s solution for 24 hours. Subsequently, sinonasal cavities were decalcified in 10% ethylenediaminetetraacetic acid (EDTA) solution at a pH of 7.4 for 10 days. After decalcification, samples were placed in a histology cassette and then paraffin embedded using the Thermo Scientific Excelsior AS tissue processor. In the processor, the tissues were embedded by undergoing the following: 2 rounds of 10% formalin for 30 minutes each, 75% alcohol for 20 minutes, 90% alcohol for 30 minutes, 95% alcohol for 30 minutes, 3 rounds of 100% alcohol for 30 minutes each, 3 rounds of xylene for 30 minutes each, 2 rounds of paraffin for 45 minutes each, and one final round of paraffin for 1 hour, respectively. The samples were then embedded in wax using the Leica EG1150H tissue embedding station. A Leica RM2255 microtome was used to section samples at 10 µm thickness. The sectioned samples were placed onto slides and stained using the periodic acid schiff (PAS) for goblet cell detection.

### Gas Chromatography - Mass Spectrometry

After collection, blood was centrifuged at 4°C for 15 minutes at 2,000 x g to allow for serum separation. Serum was transferred to a fresh tube and placed at -70°C. Serum was shipped on dry ice to the ASU-Mayo Metabolomics Lab for gas chromatography-mass spectrometry (GC-MS). For SCFA extraction, 20 µL of serum was mixed with 30 µL of 0.1 M NaOH. 20 µL of 200 µM hexanoic acid-6,6,6-d3 was added as an internal standard. To each mixture, 430 µL of MeOH was added and vortexed for ten seconds, then placed at -20°C for 20 minutes. Samples were centrifuged at 14,000 rpm for ten minutes at 4°C and 450 µL were moved to a new tube. Samples were left to dry under vacuum at 37 °C for 120 min using a CentriVap Concentrator (Labconco, Fort Scott, KS). Derivatization proceeded by adding 40 µL of methoxyamine hydrochloride solution in pyridine (MeOX, 20 mg/mL) and incubating at 60°C for 90 minutes. After incubation, 60 µL of MTBSTFA was added and incubated for an additional 30 minutes at 60°C. The sample was vortexed and centrifuged at 14,000 rpm for 10 minutes. 70 µL of supernatant was transferred to a new glass vial to be used on the Agilent 7820A gas chromatography system coupled to an Agilent 5977B mass spectrometer (Agilent Technologies, Santa Clara, CA). Separation occurred with a HP-5 ms capillary column coated with 5% phenyl-95% methylpolysiloxane (30 m×250 µm i.d., 0.25 µm film thickness, Agilent Technologies).

### 16S rRNA Gene Sequencing

The barcoded primers 515F/806R were used to target the V4 region of the 16S rRNA gene as previously described [56]. Each PCR reaction contained 2.5 µl of PCR buffer (TaKaRa, 10x concentration, 1x final), 1 µl of the Golay barcode tagged forward primer (10 µM concentration, 0.4 µM final), 1 µl of bovine serum albumin (Thermofisher, 20 mg/mL concentration, 0.56 mg/µl final), 2 µl of dNTP mix (TaKaRa, 2.5 mM concentration, 200 µM final), 0.125 µl of HotStart ExTaq (TaKaRa, 5 U/µl, 0.625 U/µl final), 1 µL reverse primer (10 µM concentration, 0.4µM final). For fecal and cecal samples, 1 µL of template DNA was included. For sinus samples, 3 µL of template DNA was included. All PCR reactions were filled to a total 25 µL with PCR grade water (Sigma-aldrich) then placed on a ThermalCycler. In short, a 30 cycle polymerase chain reaction (PCR) was used for cecum and fecal samples and a 35 cycle PCR was used for the low bacterial burden sinonasal samples. ThermalCycler conditions were as follows, 98°C denaturing step for 2 minutes, 30 or 35 cycles of 98°C for 20 seconds, 50°C for 30 seconds, and 72°C for 45 seconds, a final step of 72°C for 10 minutes. PCR was performed in triplicate for each sample and an additional negative control was included for each barcoded primer. A post-PCR quality control step was performed using a 2% agarose gel (ThermoFisher). Extraction blank controls were processed through the 16S PCR identically to tissue samples. Barcode primer NTCs controls were carried through the agarose gel step. If amplification was present for negative controls then contamination likely occurred and the PCR was repeated with a new barcoded 806R primer. Following agarose gel, PCR product was quantified using the Qubit dsDNA High Sensitivity Kit (ThermoFisher) and the Qubit fluorometer 4. PCR products were pooled at equimolar concentrations of 50 ng. Primer dimer and human mitochondrial bands, present in sinonasal samples, were removed from the final pool using a 2% E-gel size-select II (ThermoFisher). Quality of the pool was assessed with the Bioanalyzer DNA 1000 chip (Agilent Technologies) then combined with 1% PhiX. A total of 4 pools were sequenced on the Illumina MiSeq using the 600-cycle MiSeq Reagent Kit V3 (Illumina). Each pool contained 3 identical samples to assess sequencing run variations.

### Microbiome Bioinformatics

Paired-end sequences were analyzed using Quantitative Insights Into Microbial Ecology 2 (QIIME 2) version 2019.10 [57,58]. Demultiplexed sequences were denoised, trimmed at a median Q score of 30, and grouped into amplicon sequence variants (ASVs) using dada2 [59]. A phylogenetic tree was built by aligning sequences with MAFFT and a phylogenetic tree was built using FastTree2 [60,61] which was subsequently rooted by midpoint rooting. Taxonomy was assigned to ASVs with a Naive Bayes classifier trained on the Greengenes 13_8 99% OTU database [62], using the q2-feature-classifier taxonomy classification plugin [63]. ASVs corresponding to mitochondria or chloroplast, ASVs with a frequency less than 10, or ASVs observed in fewer than two samples were filtered out. The resulting table of ASV counts on a per sample basis (i.e., the ASV table) was used to compute alpha and beta diversity measures with q2-diversity. Richness (Observed ASVs) and Shannon diversity were used to measure alpha diversity (within-sample diversity). Raw alpha diversity values were imported in GraphPad PRISM version 8.0.0 (GraphPad Software, La Jolla California USA) to develop violin plots for visualization. Volatility plots were developed using q2-longitudinal to view longitudinal fecal alpha diversity [64]. Unweighted Unifrac, Weighted Unifrac, Jaccard, and Bray-Curtis were used to measure beta diversity and Principal Coordinates (PCoA) plots were visualized using Emperor [65,66]. Differential abundance was assessed using the ANCOM method in q2-composition [67]. Feature frequencies were visualized in R with ggplot 2. Machine learning classification was performed using Random Forests classifiers as implemented in q2-sample-classifier [68]. Raw feature importance was downloaded and plotted using the R libraries ggplot 2 and tidyverse. For longitudinal analysis of fecal samples, Unweighted Unifrac, Weighted Unifrac, Jaccard, and Bray-Curtis were calculated to measure beta diversity. PCoAs were built with QIIME 2 emperor to visualize fecal beta diversity results. A custom axis of day was added to the Jaccard PCoA to view supplemental changes on each collection day. To evaluate changes in the fecal microbiome on the final day of the study, corresponding to the sample time point we assessed for cecum samples, we filtered the feature table to include only samples from the final day of fecal collection. Alpha diversity and machine learning of the final day fecal samples were analyzed identically to the previously described cecum samples.

### Statistical Analysis

All statistical results were considered to be significant based on an alpha threshold of 0.05. Alpha diversity significance was tested using Kruskal-Wallis p-values generated in q2-diversity that were adjusted using the FDR method in R with the R stats package [69]. Differences in beta diversity metrics by diet or supplement were evaluated using multivariate permutational analysis of variance (PERMANOVA) using QIIME 2, and linear mixed effects models using q2-longitudinal [64]. Immune, SCFA, and goblet cell data were analyzed using paired Mann-Whitney U tests. Alpha diversity was analyzed linear mixed effects models as implemented in q2-longitudinal [64] and with Kruskal-Wallis tests as implemented in QIIME 2, and p-values were adjusted for multiple comparisons using FDR adjustments.

### Data Availability

Demultiplexed ASVs are available via the National Center for Biotechnology Information (NCBI) sequence read archive (SRA SUB7343119). Uploaded files have been demultiplexed and have relevant metadata information.

## ACKNOWLEDGEMENTS

We would like to thank the Southwest Health Equity Research Collaborative (NIH/NIMHD RCMI U54MD012388) and the Partnership of Native American Cancer Prevention (NIH/NCI U54CA143925) for funding. Thanks to the Northern Arizona University (NAU) animal facility staff for husbandry and Aubrey Funke from the NAU Imaging and Histology Core for histology. Thanks to Krystal Sheridan and Heather Centner with the Translational Genetics and Genomics North for their role in the Joint Sequencing Core. Thanks to Tate and Lyle for providing the fiber. SHERC and Tate and Lyle had no role in the study design, data collection, nor analysis for this study.

