## Supplemental Figures for "Soluble corn fiber reduces ovalbumin-induced sinonasal inflammation via the gut microbiota-airway axis"

SUPPLEMENTAL INFORMATION

**TABLE S1** Cellulose supplemented ingredient matched diets[a]

| Formula | Control Diet + Cellulose (g/kg) | Western Diet + Cellulose (g/kg) |
|---|---|---|
| Casein | 190 | 236 |
| DL-Methionine | 3 | 3.54 |
| Sucrose | 50 | 179.12 |
| Corn Starch | 485.79 | 145.8 |
| Maltodextrin | 120 | 120 |
| Anhydrous Milkfat | 19.5 | 100 |
| Beef Tallow | 19.5 | 100 |
| Soybean Oil | 11 | 3.5 |
| Cellulose | 52.2 | 54.2 |
| Mineral Mix | 35 | 41.3 |
| Calcium Phosphate | 4 | 4.72 |
| Vitamin Mix | 10 | 11.8 |
| Ethoxyquin | 0.01 | 0.02 |

[a]Insoluble cellulose was supplemented directly into the rodent chow.

**TABLE S2** RT-PCR Gene targets[a]

| Assay Targets | $T_H1$ Marker | $T_H2$ Marker |
|---|---|---|
| Arginase | | X |
| Chemokine (C-C motif) ligand 11 | | X |
| Interferon gamma | X | |
| Interleukin 1 beta | X | |
| Interleukin 4 | | X |
| Interleukin 5 | | X |
| Interleukin 6 | X | |
| Interleukin 13 | | X |
| Interleukin 17A | | |
| Mucin 5, subtypes A and C, tracheobronchial/gastric | | X |
| Nitric oxide synthase 2, inducible | | X |
| Transforming growth factor, beta 1 | X | |
| Glyceraldehyde-3-phosphate dehydrogenase | | |
| Genomic DNA Control | | |
| Positive PCR Control | | |
| Reverse Transcription Control | | |

[a]A custom qPCR array was developed to screen different $T_H1$ and $T_H2$ gene markers. An X represents the category the gene is considered.

**TABLE S3** PERMANOVA Beta diversity results[a]

| Distance Metric | Sample Type | Categorical Variable | pseudo-F | PERMANOVA p value |
|---|---|---|---|---|
| **Weighted Unifrac** | **Cecum** | **diet** | **7.67917** | **0.001** |
| **Unweighted Unifrac** | **Cecum** | **diet** | **6.2809** | **0.001** |
| **Jaccard** | **Cecum** | **diet** | **3.80736** | **0.001** |
| **Bray-Curtis** | **Cecum** | **diet** | **6.24022** | **0.001** |
| **Weighted Unifrac** | **Cecum** | **supplement** | **4.41743** | **0.002** |
| **Unweighted Unifrac** | **Cecum** | **supplement** | **6.62723** | **0.001** |
| **Jaccard** | **Cecum** | **supplement** | **5.7161** | **0.001** |
| **Bray-Curtis** | **Cecum** | **supplement** | **8.64515** | **0.001** |
| Weighted Unifrac | Cecum-SCF only | diet | 1.9663 | 0.12 |
| **Unweighted Unifrac** | **Cecum-SCF only** | **diet** | **3.58663** | **0.005** |
| **Jaccard** | **Cecum-SCF only** | **diet** | **2.38138** | **0.017** |
| Bray-Curtis | Cecum-SCF only | diet | 2.09405 | 0.085 |
| Weighted Unifrac | Final Day Fecal | diet | 1.86445 | 0.133 |
| **Unweighted Unifrac** | **Final Day Fecal** | **diet** | **3.81983** | **0.001** |
| **Jaccard** | **Final Day Fecal** | **diet** | **2.82657** | **0.002** |
| **Bray-Curtis** | **Final Day Fecal** | **diet** | **2.65247** | **0.024** |
| **Weighted Unifrac** | **Final Day Fecal** | **supplement** | **3.22375** | **0.007** |
| **Unweighted Unifrac** | **Final Day Fecal** | **supplement** | **6.37449** | **0.001** |
| **Jaccard** | **Final Day Fecal** | **supplement** | **5.26479** | **0.001** |
| **Bray-Curtis** | **Final Day Fecal** | **supplement** | **4.75649** | **0.001** |
| Weighted Unifrac | Sinonasal cavity | diet | 2.01434 | 0.078 |
| Unweighted Unifrac | Sinonasal cavity | diet | 1.2005 | 0.201 |
| Jaccard | Sinonasal cavity | diet | 1.10905 | 0.226 |
| Bray-Curtis | Sinonasal cavity | diet | 1.37179 | 0.158 |
| Weighted Unifrac | Sinonasal cavity | supplement | 1.41403 | 0.195 |
| Unweighted Unifrac | Sinonasal cavity | supplement | 1.04296 | 0.342 |
| Jaccard | Sinonasal cavity | supplement | 1.08858 | 0.196 |
| Bray-Curtis | Sinonasal cavity | supplement | 1.38627 | 0.085 |

[a]PERMANOVA p-values of various beta diversity metrics on various sample types.

**Fig. S1**

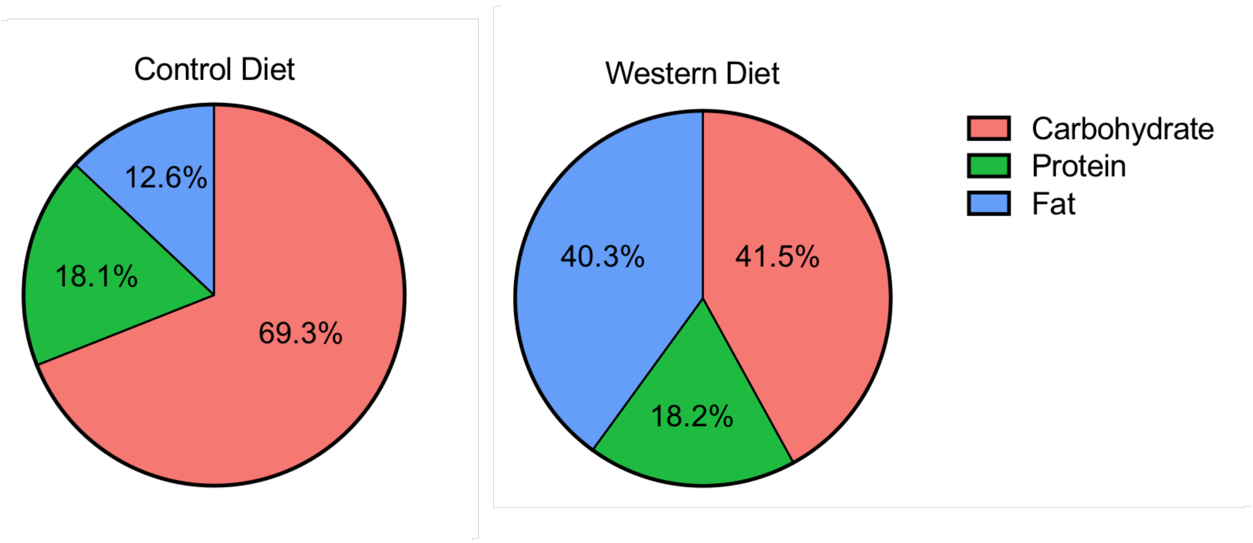


**Figure S1:** Macronutrient breakdown of murine diets.

**Fig. S2**

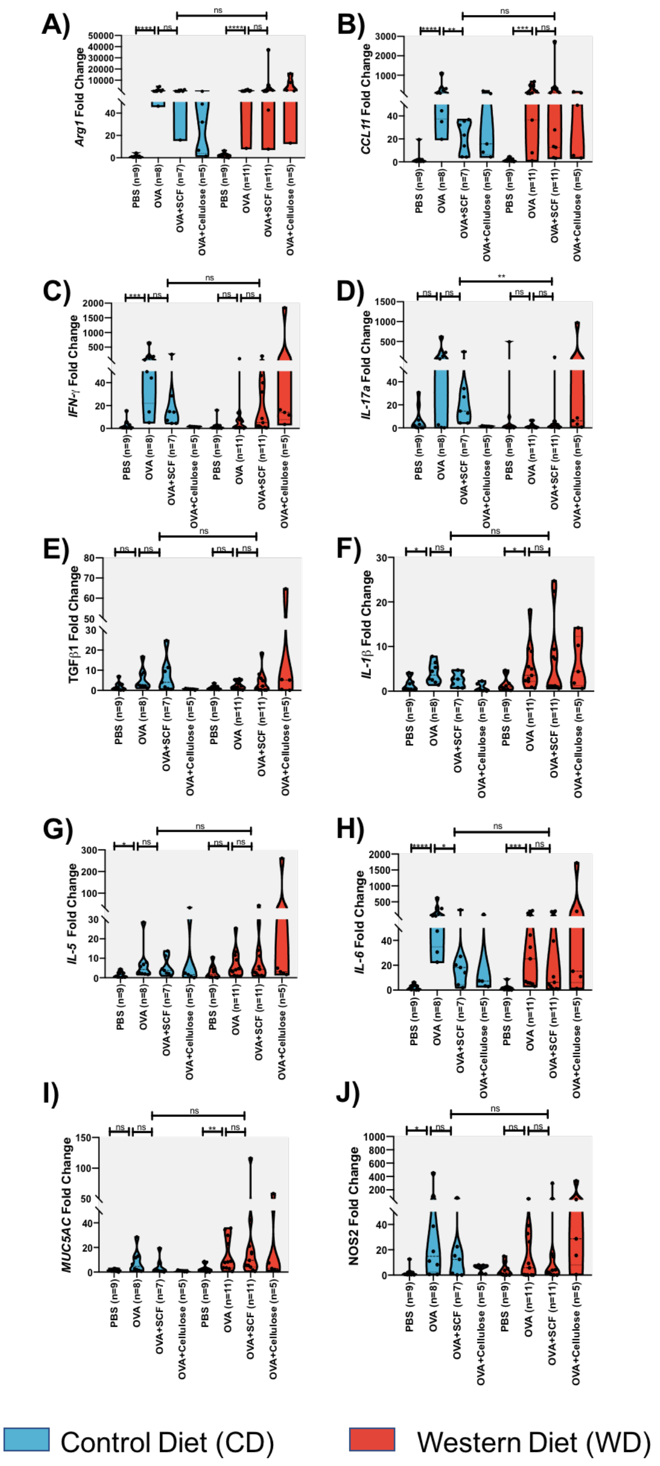


**Figure S2:** Sinonasal cavity gene expression results of all tested $T_H1$ and $T_H2$ genes.

Mice underwent allergic airway sensitization with ovalbumin (OVA) or received a vehicle control (PBS). A subset of mice were fed a soluble corn fiber supplement (SCF). Mice were maintained on either a control diet (CD) or western diet (WD). Reverse transciptase qPCR was used to measure $T_H1$ and $T_H2$ inflammation markers in the sinonasal cavities. **A)** Arginase 1 (*Arg1*) **B)** Chemokine ligand 11 (*CCL11*) **C)** Interferon gamma (*IFN-γ*) **D)** Interleukin 17A (*IL-17A*) **E)** Transforming growth factor beta 1 (*TGFβ1*) **F)** Interleukin 1 beta (*IL-1β*) **G)** Interleukin 5 (*IL-5*) **H)** Interleukin 6 (*IL-6*) **I)** Mucin 5, subtypes A and C, tracheobronchial/gastric (*MUC5AC*) **J)** Nitric oxide synthase 2, inducible (*NOS2*).

**Fig. S3**

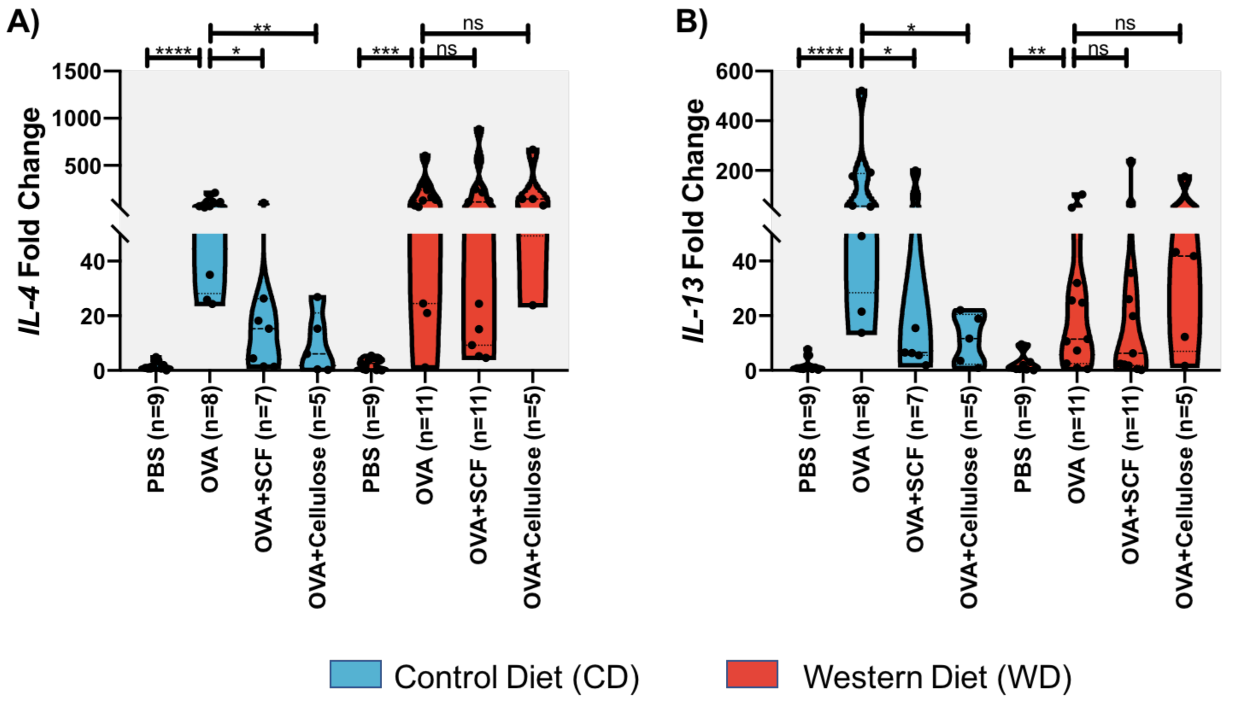


**Figure S3**: Gene expression of type-2 inflammation markers in mouse sinonasal cavities. Mice underwent allergic airway sensitization with ovalbumin (OVA) or received a vehicle control (PBS). Mice were maintained on either a control diet (CD) or western diet (WD). Mice either received no supplement, a soluble corn fiber (SCF) supplement, or an insoluble fiber (Cellulose) supplement. qPCR was used to measure $T_H2$ inflammation markers in the sinonasal cavities. **A)** *IL-4*, a marker of $T_H2$ inflammation was reduced in CD-fed cellulose treated mice (p=0.0031, Mann Whitney) **B)** *IL-13*, a second marker of $T_H2$ inflammation was reduced in cellulose treated mice consuming a control diet (p=0.0031, Mann Whitney).

**Fig. S4**

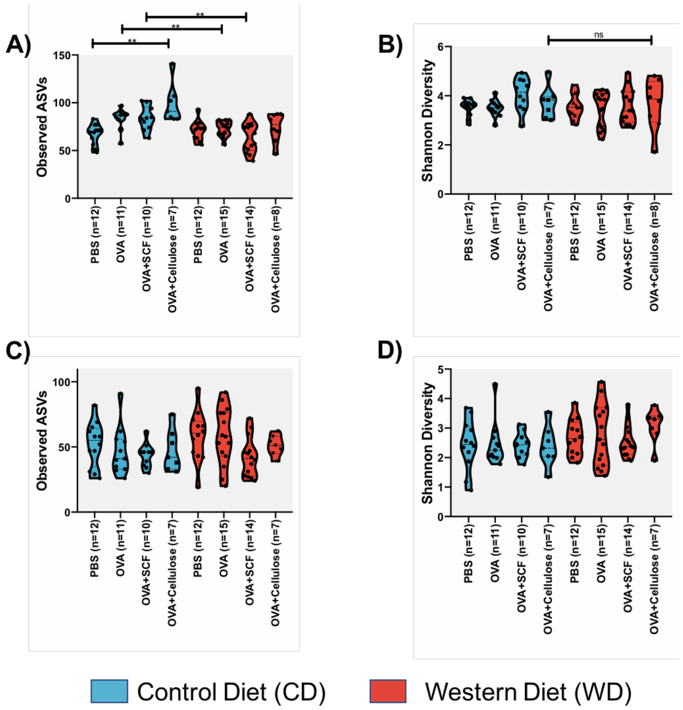


**Figure S4:** Alpha diversity of cecum samples. Mice underwent allergic airway sensitization with ovalbumin (OVA) or received a vehicle control (PBS). Mice were maintained on either a control diet (CD) or western diet (WD). Mice either received no supplement, a soluble corn fiber (SCF) supplement, or an insoluble fiber (Cellulose) supplement. Amplicon sequencing of the V4 16S rRNA gene region was performed and used to measure alpha diversity differences of cecum and fecal samples. **A)** Observed amplicon sequencing variants (ASVs) are greater in CD-fed and cellulose treated cecal samples than in WD-fed and cellulose treated mice, however it is not significant (p=0.069, Kruskal-Wallis, FDR adjusted p-values) **B)** There are no differences in Shannon diversity of cecal samples (Kruskal-Wallis, FDR adjusted p-values) **C)** Alpha diversity of fecal samples using the observed ASVs metric **D)** Alpha diversity of fecal samples using the Shannon metric (Kruskal-Wallis, FDR adjusted p-values).

**Fig. S5**

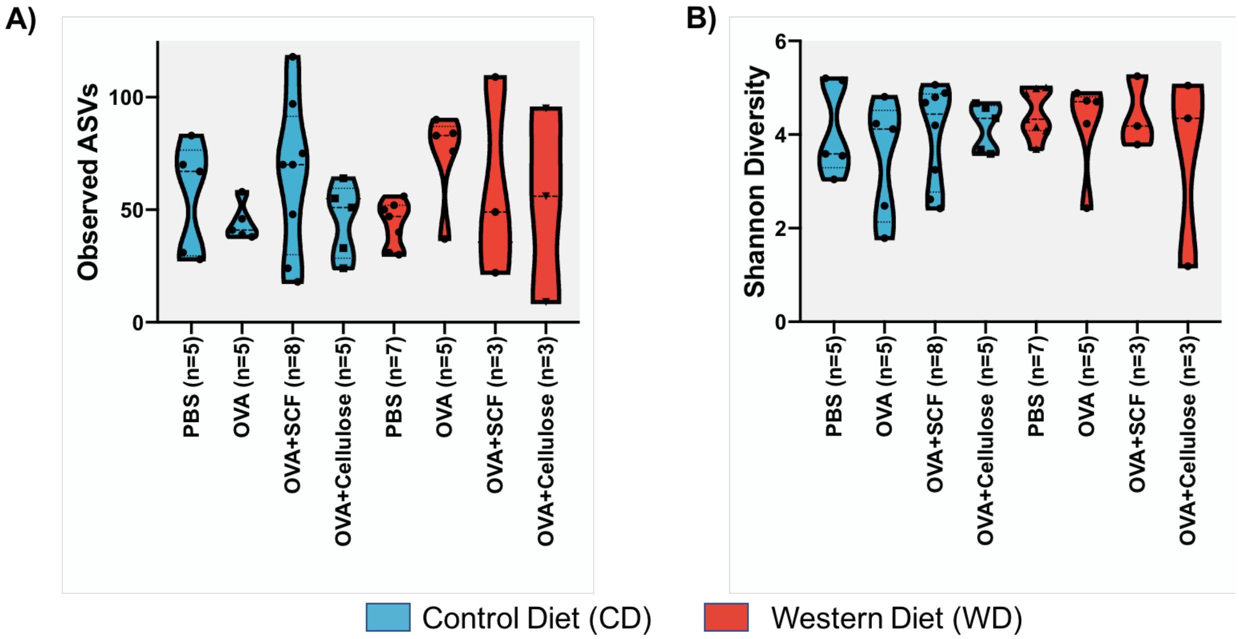


**Figure S5**: There are no differences in alpha diversity of sinonasal samples. Mice underwent allergic airway sensitization with ovalbumin (OVA) or received a vehicle control (PBS). Mice were maintained on either a control diet (CD) or western diet (WD). Mice either received no supplement, a soluble corn fiber (SCF) supplement, or an insoluble fiber (Cellulose) supplement. Amplicon sequencing of the V4 16S rRNA gene region was performed and used to measure alpha diversity differences of the sinonasal cavity samples. **A)** Observed ASVs in the sinonasal microbiome **B)** Shannon diversity in the sinonasal microbiome.

**Fig. S6**

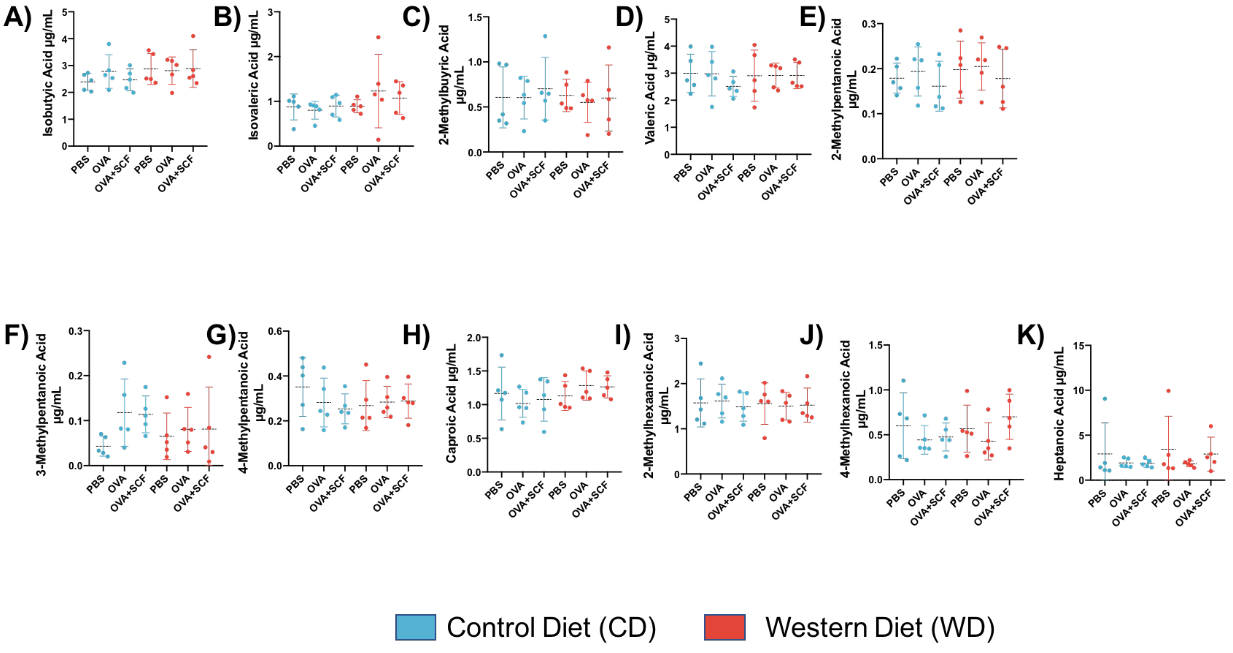


**Figure S6:** Mice were maintained on a control diet (CD) or western diet (WD) with either no supplement, a soluble corn fiber (SCF) supplement, or an insoluble fiber (Cellulose). Serum was collected on the final collection day. Gas chromatography-mass spectrometry was used to quantify short-chain fatty acids in serum samples. (CD: control diet, WD: western diet, PBS : no allergic airway sensitization, OVA : allergic airway sensitization with ovalbumin, OVA + SCF : allergic airway sensitization and a soluble corn fiber supplement). **A)** Isobutyric acid **B)** Isovaleric Acid **C)** 2-Methylbutyric acid **D)** Valeric acid **E)** 2-Methylheptanoic acid **F)** 2-Methylpentanoic acid **G)** 4-Methylpentanoic acid **H)** Caproic acid **I)** 2-Methylhexanoic acid **J)** 4-Methylhexanoic acid **K)** Heptanoic acid.

**Fig. S7**

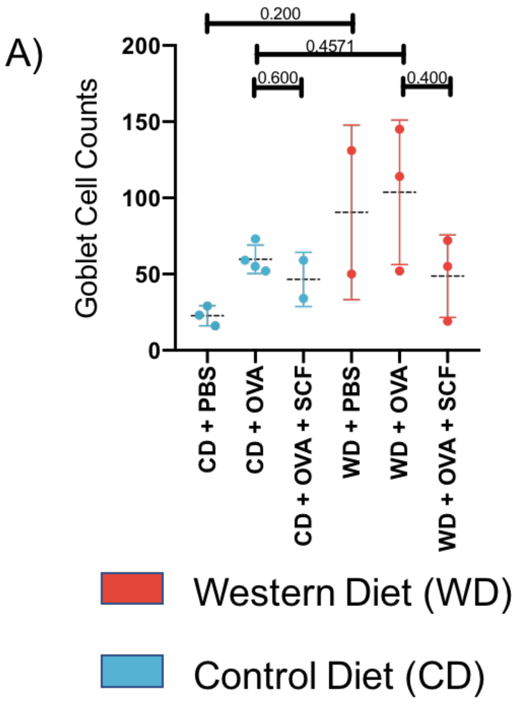


**Figure S7:** Mice underwent allergic airway sensitization with ovalbumin (OVA) or received a vehicle control (PBS). Mice were maintained on either a control diet (CD) or western diet (WD). Mice either received no supplement, a soluble corn fiber (SCF) supplement, or an insoluble fiber (Cellulose) supplement. Sinonasal cavities were collected on the last day of the experiment and goblet cells were stained using the periodic-acid shiff (PAS) stain. No significant differences between groups were identified (Mann-Whitney $p>0.05$).
